# Multivalent interactions facilitate motor-dependent protein accumulation at growing microtubule plus ends

**DOI:** 10.1101/2021.09.14.460284

**Authors:** Renu Maan, Louis Reese, Vladimir A. Volkov, Matthew R. King, Eli van der Sluis, Nemo Andrea, Wiel Evers, Arjen J. Jakobi, Marileen Dogterom

**Author notes:** These authors contributed equally (Renu Maan, Louis Reese, Vladimir A. Volkov).

## Abstract

Growing microtubule ends provide platforms for the accumulation of plus-end tracking proteins that organize into comets of mixed protein composition. Using a reconstituted fission yeast system consisting of end-binding protein Mal3, kinesin Tea2 and cargo Tip1, we found that these proteins can be driven into liquid phase droplets both in solution and at microtubule ends under crowding conditions. In the absence of crowding agents, cryo-electron tomography revealed that motor-dependent comets consist of disordered networks where multivalent interactions appear to facilitate the non-stoichiometric accumulation of cargo Tip1. We dissected the contribution of two disordered protein regions in Mal3 and found that both are required for the ability to form droplets and Tip1 accumulation, while autonomous Mal3 comet formation only requires one of them. Using theoretical modeling, we explore possible mechanisms by which motor activity and multivalent interactions may lead to the observed enrichment of Tip1 at microtubule ends.

## Introduction

Growing microtubule plus ends recruit an evolutionary conserved network of proteins interacting with so-called end-binding (EB) proteins (Honnappa et al., 2009). This network exists as a multivalent protein assembly which recognizes features of growing microtubule ends, such as GTP hydrolysis intermediates (Maurer et al., 2014), bent tubulin protofilaments (Guesdon et al., 2016), and tubulin interfaces which are unavailable on closed microtubules (Reid et al., 2019).

In fission yeast *Schizosaccharomyces pombe* and filamentous fungi *Aspergllus nidulans*, the microtubule plus end tracking system is crucial to establish cell polarity by asymmetrically transporting and distributing polarity markers to the cellular cortex (Browning et al., 2003; Brunner and Nurse, 2000). Once associated with the cellular cortex, many of these markers behave like multi-protein complexes or clusters (Dodgson et al., 2013), which raises the question how proteins interact with each other at growing microtubule ends and whether clusters may already be formed before being deposited at the cortex (Taberner and Dogterom, 2019).

A minimal protein network for <u>m</u>icrotubule <u>p</u>lus <u>e</u>nd tracking (MPET) was first reconstituted *in vitro* using purified proteins from *S. Pombe* (Bieling et al., 2007a). The three proteins that are necessary and sufficient for successful *in vitro* plus-end tracking are Mal3 (EB homolog), Tea2 (kinesin-7 homolog) and Tip1 (CLIP-170 homolog). Accumulation of Tip1 and Tea2 at the microtubule end is Mal3-dependent both *in vitro* and *in vivo* (Browning et al., 2003; Busch and Brunner, 2004). Mal3 is needed for ATPase activity and processive transport of Tea2 (Browning and Hackney, 2005). However, affinity of Mal3 for microtubules is independent of Tea2 and Tip1. Tip1 has been shown to interact with the EB homology domain of Mal3 through its CAP-Gly domain (Busch and Brunner, 2004), as also shown for Tip1 homolog CLIP170 and other plus-end tracking proteins (+TIPs) interacting with EB proteins (Bieling et al., 2008; Hayashi et al., 2005; Honnappa et al., 2009; Stangier et al., 2018). Tea2 interacts with Mal3 through its N-terminal extension and with Tip1 through its coiled-coil region (Browning and Hackney, 2005; Brunner and Nurse, 2000; Busch et al., 2004). Since many of these interactions happen through unstructured protein regions (Supplementary Fig. S1A), we hypothesize that the Mal3/Tip1/Tea2 network may be formed by multivalent low-affinity interactions that are a hallmark of liquid-liquid phase separation (LLPS) (Alberti et al., 2019; Wang et al., 2018).

LLPS is the phenomenon of reversible de-mixing of miscible components from their homogeneous mixture driven by microscopic interactions between the molecules (Hyman et al., 2014). Eukaryotic cells contain many membrane-bound and membrane-less organelles which form through similar phase separation processes. Examples include Cajal bodies, nuclear speckles, nucleolus, stress granules and P-bodies (Banani et al., 2017; Brangwynne et al., 2009; Woodruff et al., 2015). Recently a number of microtubule associated proteins (MAPs) have been reported to undergo similar demixing *in vitro* with proposed relevance for microtubule dynamics, nucleation, branching etc. (Hernández-Vega et al., 2017; King and Petry, 2020; Setru et al., 2021; Siahaan et al., 2019; Tan et al., 2019). While the importance of these liquid- and gel-like assemblies for cellular function is still controversial (Alberti et al., 2019; McSwiggen et al., 2019; Raff, 2019), it is widely accepted that disordered protein regions often drive interactions leading to phase separation (Wang et al., 2018).

Here we investigate the role of multivalent interactions in the formation of comets at growing microtubule ends in an *in vitro* reconstitution experiment. Under crowding conditions, we observed the formation of phase separated droplets of fission yeast MPET proteins, both in the absence and presence of microtubules. Driven by motor activity, these droplets fuse into larger droplets at growing microtubule ends. Cryogenic electron tomography (cryoET) revealed that the comets formed at microtubule plus ends by Mal3, Tea2 and Tip1 in the absence of crowding agents appear as a loose disordered network. These networks are distinct from dense PEG-driven droplets but may nevertheless facilitate the non-stoichiometric accumulation of Tip1 at growing microtubule ends, as observed by fluorescence microscopy. Starting with these observations we dissect contributions of molecular interactions via intrinsically disordered regions (IDRs) to both the formation of microtubule end-tracking comets, and the ability of Mal3/Tip1/Tea2 to phase separate in crowding conditions. Using truncated constructs of Mal3, we show that deletion of Mal3 IDR1 disrupts both comet formation and LLPS of Mal3. Removal of Mal3 IDR2 also disrupts LLPS of Mal3 as well as motor-driven end-accumulation of Tip1, but it does not affect the autonomous ability of Mal3 to form comets. We conclude that multivalent interactions contribute to the network-like architecture of plus end comets, forming disordered structures that are easily driven into phase-separated dense droplets under crowding conditions. We propose that these non-stoichiometric structures allow for the efficient motor-driven accumulation of Tip1 at microtubule ends.

Finally, we use stochastic modelling to understand how motor transport and multivalent interactions may contribute to the efficient formation of Tip1 comets. In agreement with our observations by fluorescence microscopy and cryo-ET, the modelling shows that motor-based transport, multivalent interactions between the components, and the presence of a distinct microtubule end structure may collectively contribute to the formation of dense Tip1 comets at microtubule ends.

## Results

### Plus-end trackers form a multi-protein complex on the microtubule lattice and ends

We reconstituted the fission yeast MPET network *in vitro* using bacterially expressed proteins Mal3, Tea2, and Tip1, as reported previously (Bieling et al., 2007a) (**Fig. 1A**). Using total internal reflection fluorescence (TIRF) microscopy and double labelling of either Mal3-Alexa647 and Tip1:GFP (**Fig. 1B**), or Mal3-Alexa488 and Tea2-Alexa647 (**Fig. 1C**), we observed that all three proteins were transported on the microtubule lattice in the direction of the microtubule plus end and were all present in an end-tracking comet. Therefore, the three proteins in the fission yeast MPET network co-exist in a multiprotein complex when interacting with the microtubule lattice. Since all three proteins contain disordered, low-complexity regions (**Supplementary Fig. S1A**), we hypothesized that efficient plus-end accumulation of the Mal3/Tea2/Tip1 protein network is facilitated through multivalent or non-stochiometric protein interactions. To test this hypothesis, we investigated the behavior of the protein network under crowding conditions.

**Figure 1:**
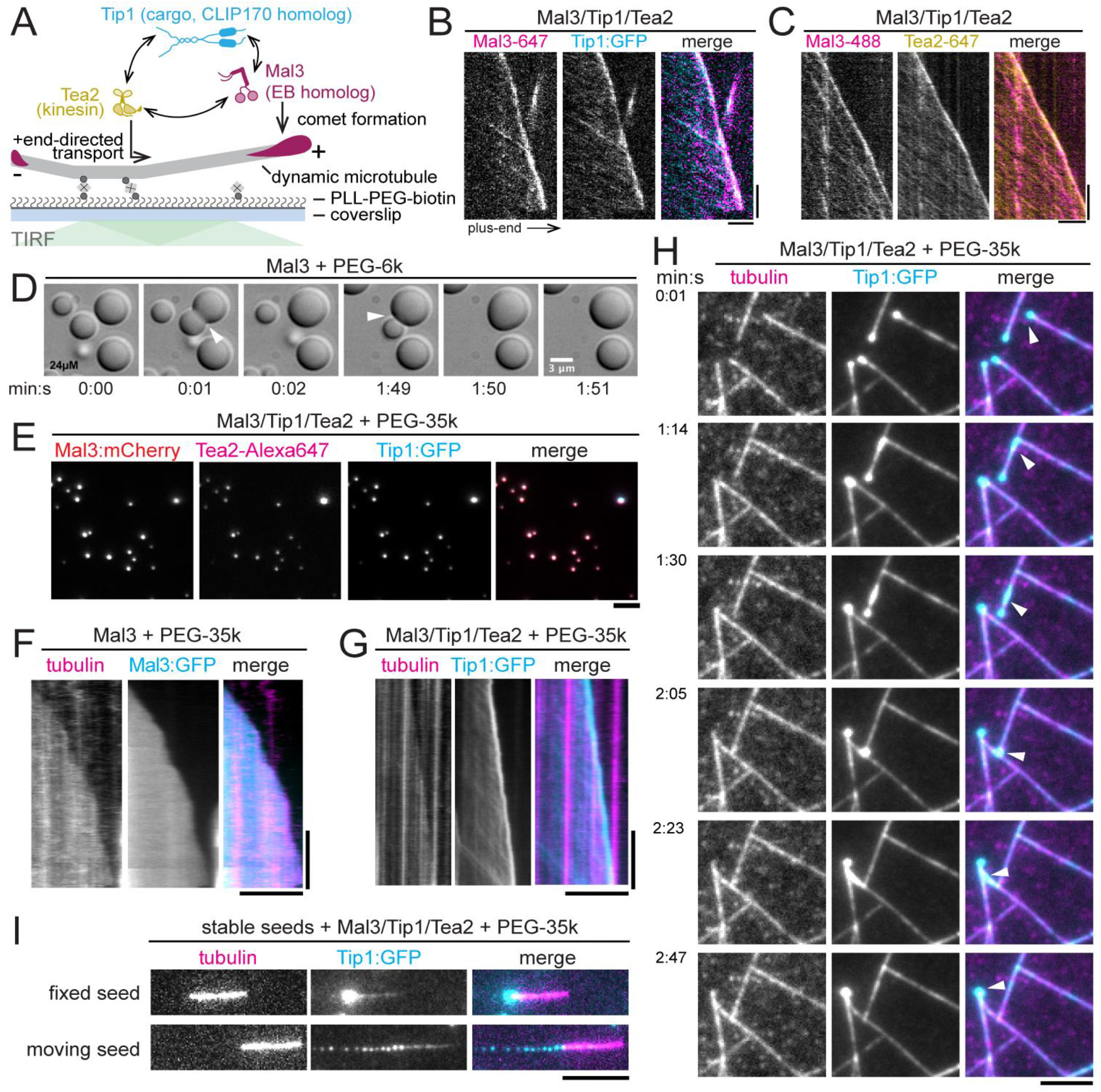
Fission yeast microtubule end-tracking system undergoes phase separation under crowding conditions *in vitro*. **(A)** Cartoon showing the interactions among the three plus end tracking proteins, Mal3, Tea2 and Tip1. **(B & C)** Kymographs of MPET (microtubule plus-end tracking) reconstitutions showing co-localization of Mal3-Alexa647 and Tip:GFP (B), and Mal3-Alexa488 and Tea2-Alexa647 (C) on both the microtubule lattice and the plus end. Scale bar: 5 µm and 60 sec. **(D)** Large Mal3 condensates form in the presence of 10% (w/v) PEG 6k at 24 µM concentration. In the panel two condensates can be seen fusing over time (arrowheads). **(E)** Co-condensation of Mal3:mCherry (200 nM), Tea2-Alexa647 (20 nM) and Tip1:GFP (150 nM) in the presence of 5% PEG 35k at the protein concentrations used in the MPET reconstitution assays. **(F)** Mal3:GFP (200 nM) coats the entire microtubule lattice in the presence of 5% PEG 35k while there is no distinct accumulation at the microtubule end. **(G)** Combination of Mal3 (200 nM) with Tea2 (20 nM) and Tip1:GTP (150 nM) in the presence of 5% PEG 35k leads to Tip1:GFP accumulation at the plus end with motor traces visible on the lattice. **(H)** A droplet of Mal3/Tea2/Tip1 formed at the plus end of one microtubule gets transferred to the lattice of another (arrowhead). The transferred droplet spreads on the microtubule lattice and moves toward the plus end where it fuses with the already existing Mal3/Tea2/Tip1 droplet. See also Video S2. Scale bars: 5 µm. **(I)** Top panel shows MPET reconstitution 4on GMPCPP stabilized seeds in the presence of 5% PEG 35k. Bottom panel shows deposition of Mal3/Tea2/Tip1 droplets by the moving seed on the glass surface in the presence of 5% Peg 35k. Seed movement occurs through non-specific binding of Tea2 to the surface.

### Mal3, Tea2, and Tip1 form co-condensates under crowding conditions

We performed reconstitution assays under crowding conditions both without and with microtubules. We first mixed all MPET network components and studied their behavior in solution. To study how dynamic microtubules affect these interactions, we next performed reconstitutions on functionalized glass surfaces in the presence of GMPCPP-stabilized microtubule seeds and soluble tubulin.

Since Mal3 is an autonomous end-tracker and also plays a key role in motor activation needed for plus end tracking of the MPET network (Bieling et al., 2007a), we first focused on the ability of Mal3 to form condensates, its end-tracking ability, and its interactions with Tea2 and Tip1. At high concentrations, Mal3 readily formed condensates in the presence of PEG 6k (**Fig. 1D**) which fused together like fluid droplets (**Video S1**). To probe the robustness of droplet formation, we systematically explored the effects of Mal3 and PEG concentration as well as PEG chain length. At 200 nM Mal3:mCherry, a typical concentration in microtubule end-tracking assays, and 5% (w/v) of PEG 35k, Mal3 produced robust protein droplets (**Supplementary Fig. S1B**). In the additional presence of 20 nM Tea2-Alexa647 and 150 nM Tip1:GFP, typical concentrations for microtubule end-tracking reconstitutions, we observed co-localization of Tea2 and Tip1 with Mal3 condensates (**Fig. 1E**). Also, Tea2 and Tip1 formed condensates under similar crowding conditions and concentrations on their own (**Supplementary Fig. S4**).

### Co-condensation of Mal3, Tea2, and Tip1 in the presence of microtubules

To establish how crowding-induced droplets interact with microtubules, three different assays were designed. We reconstituted MPET in the presence of crowding agent PEG 35k on static and dynamic microtubules attached to functionalized glass surfaces and used GMPCPP-stabilized microtubule seeds with and without soluble tubulin and GTP.

In the first assay, microtubules were grown from biotinylated seeds attached to the coverslip in a flow cell. In the presence of crowding agent, Mal3:GFP coated the entire microtubule lattice and no difference was found between the fluorescence on the plus end and the lattice (**Fig. 1F**). When Tea2 and Tip1:GFP were added to the PEG-containing assay, we observed both motor traces at the lattice and bright comets at microtubule plus-ends (**Fig. 1G**). These plus-end-bound comets could transfer from one microtubule to another whenever a microtubule plus-end encountered another microtubule lattice (**Fig. 1H**). A comet that was transferred from the plus end of one microtubule onto the lattice of another microtubule spread out over the new lattice in a fluid-like manner and was transported again towards the plus-end. The transferred droplet/comet was then observed to merge with the already existing comet of the 2^nd^ microtubule (**Fig. 1H** and **Video S2**).

In the second assay, we added the condensates to the flow chambers with immobilized seeds on the cover glass. In the presence of PEG we observed Mal3 binding to the GMPCPP seeds, contrary to non-crowding conditions where Mal3 did not interact with the seeds (**Supplementary Fig. S1C**). In the presence of all MPET proteins we observed Tip1:GFP transport towards the plus-end on seeds. Presumably, PEG-assisted Mal3 binding to the seeds was sufficient to induce Tea2 activity and hence Tip1:GFP transport towards the plus end. Droplets were observed to form at the plus ends of the seeds that grew over time due to continuous Tea2-driven transport along the seeds (**Fig. 1I** (top panel) and **Video S3**).

In the third assay, we flushed in condensates together with seeds in the empty flow chamber. Here again we observed droplets containing Tip1:GFP being formed at the plus end of the seeds. But since the seeds were free to move and motors were non-specifically binding to the glass surface, seeds started gliding and depositing trails of droplets behind their plus-ends (**Fig. 1I** (bottom panel) and **Video S4**), similar to the Plateau-Rayleigh instability (Plateau, 1873; Rayleigh, 1878).

Together these observations provide evidence that in the presence of crowding agent Mal3, Tip1, and Tea2 together form condensates both in the absence and in the presence of microtubules. The observed condensates are liquid-like in nature, can coat the microtubule lattice, and can be transported by Tea2 motors towards the plus-ends of microtubules.

### CryoET of Mal3/Tip1/Tea2 droplets and networks at microtubule ends

The observation of droplet-like comets under crowding conditions prompted us to ask whether Ma3/Tip1/Tea2 comets are also formed by LLPS in the absence of crowding agents. Given the size of the comets in the absence of crowding agents and the limited resolution of fluorescence microscopy, we imaged Mal3/Tip1/Tea2 assemblies using cryoET.

We first added pre-formed droplets made by incubating the Mal3/Tip1/Tea2 mixture with 10% PEG 6K to holey carbon grids and vitrified them (**Fig. 2A**). In these conditions, we observed spherical droplets with fine internal grain, both in single 2D projections (**Fig. 2B**) and after 3D reconstruction of tomographic volumes (**Fig. 2C**).

**Figure 2:**
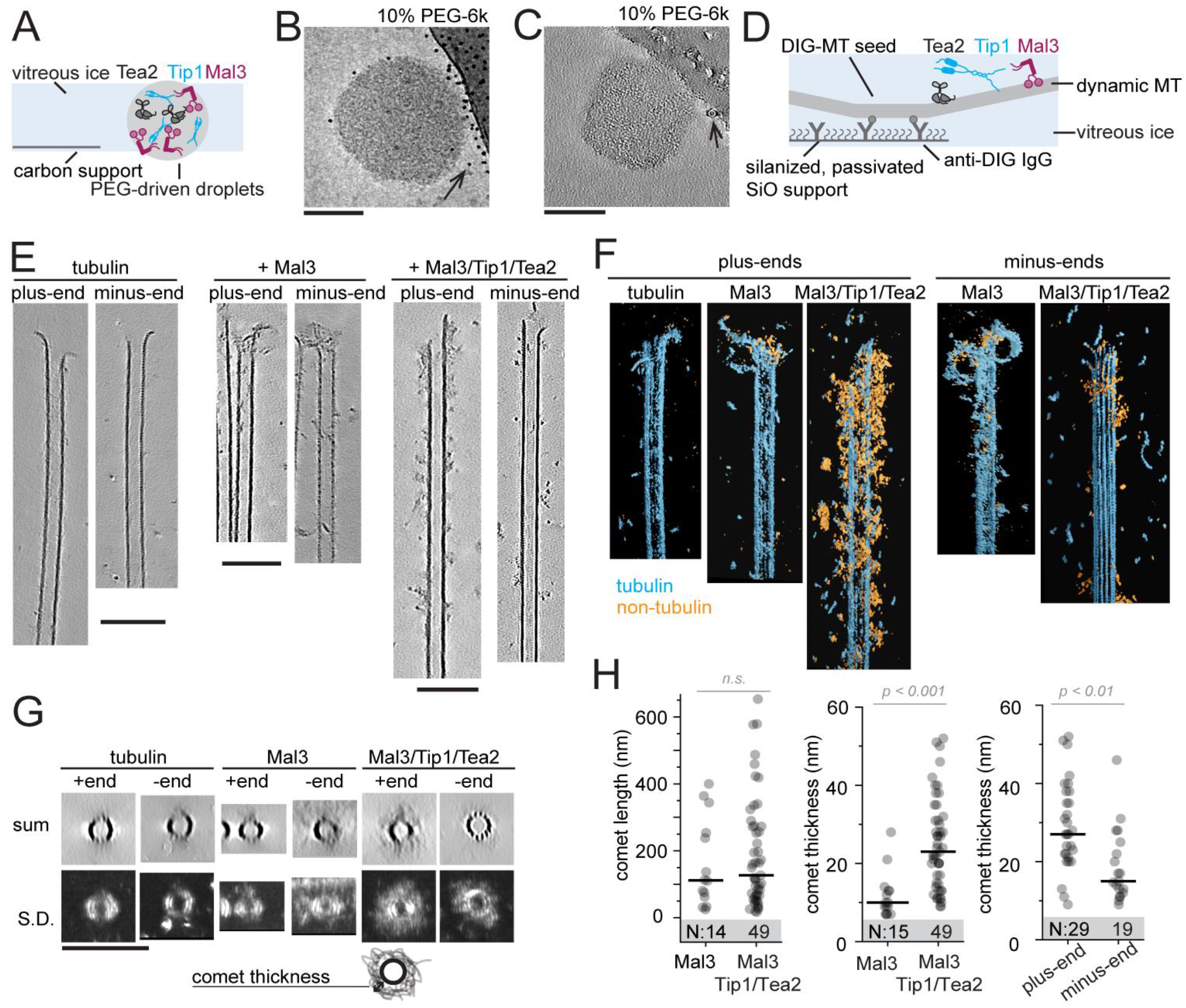
CryoET analysis of Mal3/Tip1/Tea2 assemblies. **(A)** Schematics of sample preparation for analysis of PEG-driven droplets. **(B)** A single 10-s exposure of a droplet in a hole attached sideways to a carbon support. **(C)** A 0.7 nm-thick slice through a 3D tomographic volume. Arrows in B and C show 5 nm gold beads added as fiducials for 3D reconstruction (note that the gold density is erased in the 3D volume, but not in the 2D image). **(D)** Schematics of sample preparation to study microtubule-bound assemblies of Mal3/Tip1/Tea2. **(E)** 0.7 nm-thick slices through 3D tomographic volumes recorded in the conditions indicated. (**F**) 3D renders of tomograms containing microtubule ends with bound material, segmented using *tomoseg* module of EMAN2.2 (see Materials and Methods for details). Cyan shows tubulin and microtubules, orange – all non-tubulin densities. **(G)** Projections along the microtubule length of volumes presented as sum of slices (top) and standard deviation (bottom). **(H)** Comet length (left) and thickness (right) in the presence of Mal3 alone or Mal3/Tip1/Tea2. Each datapoint represents a single microtubule end. Lines show median, numbers in the shaded area show N. Scale bars: 100 nm.

Repeating this sample preparation in the presence of dynamic microtubules, but now in the *absence* of PEG resulted in the end-tracking protein network aggregating at the carbon support (data not shown). To prevent non-specific adsorption, we adapted passivation methods previously established for treatment of glass coverslips (Volkov et al., 2018). We silanized a SiO film on the grids, adsorbed anti-DIG IgG to the silanized surface, and then made the film hydrophilic by incubation with Pluronic F-127 (**Fig. 2D**). This treatment allowed us to firmly attach GMPCPP-stabilized, DIG-labeled microtubule seeds, while rejecting the binding of other proteins from solution. We then added tubulin in the presence or absence of Mal3 alone or the complete Mal3/Tip1/Tea2 network and plunge-froze the grids after 5-7 minutes of microtubule growth.

Since we used GMPCPP seeds to nucleate microtubule growth, most microtubules in our samples contained 14 protofilament lattices (75%, see Supplementary Table S1). Characteristic moiré patterns of 14-pf microtubules made it easy to identify the polarity of their ends (**Supplementary Fig. S2A**) (Chrétien et al., 1996). In the absence of end-tracking proteins, we observed microtubules growing with flared protofilaments at their ends, as described previously (McIntosh et al., 2018), and no lattice or end decoration (**Fig. 2E** left). Adding Mal3 alone did not produce clearly visible densities at microtubule ends (**Fig. 2E** middle), but we observed a clear diffuse coating at the ends of growing microtubules when all three components Mal3, Tip1 and Tea2 were present (**Fig. 2E** right and **Supplementary Fig. S2B**).

To assist the interpretation of the reconstructed tomograms, we used volume segmentation to highlight tubulin and microtubules (cyan) and non-tubulin densities (yellow) (**Fig. 2F**). Together with polarity assignment, this allowed us to visualize massive microtubule end-bound structures at the plus ends in the presence of all three proteins Mal3, Tea2, and Tip1 (**Fig 2F, Supplementary Fig. S2B, Video S5**). Structures binding to minus ends in the presence of Mal3/Tea2/Tip1 (**Video S6**), to plus ends in the presence of Mal3 alone (**Supplementary Fig. S2C, Video S7**), or to plus ends in the absence of additional proteins (**Supplementary Fig. S2D, Video S8**) appeared much smaller. We further analyzed microtubule cross-sections to obtain quantitative information on the microtubule end-bound structures (**Fig. 2G,H**). The average thickness of comets extending outwards from the microtubule surface in the presence of Mal3 alone was 11 ± 6 nm (n = 15, here and onwards mean ± SD), considerably thinner than 29 ± 12 nm (n = 28, p < 10^−5^) in the presence of Mal3, Tip1 and Tea2 (**Fig. 2H** middle). The difference in comet length was not significant: 162 ± 132 nm for Mal3 (n = 14) and 257 ± 182 nm for Mal3/Tip1/Tea2 (n = 27, p = 0.09) (**Fig. 2H** left). Plus ends carried thicker comets of Mal3/Tip1/Tea2 (32 ± 10 nm) compared to minus ends in the same sample (21 ± 11 nm, p = 0.02) (**Fig. 2H** right). The polarity-dependent thickness of comets is consistent with the plus end-directed motility of Tea2 bringing its cargo, Tip1, to the plus ends of microtubules.

There is clearly a difference between the dense internal organization of PEG-driven droplets shown in Fig. 2B,C and the much more loosely structured microtubule-bound comets. Yet, it is possible that multivalent interactions responsible for LLPS under crowding conditions are also facilitating the formation of the network-like architecture of motor-driven plus end comets observed in cryo-EM in the absence of crowding conditions.

### Non-stoichiometric Tip1 accumulation at microtubule ends

The network-like architecture described above may facilitate the non-stoichiometric accumulation of Tip1 cargo at microtubule ends as also observed by fluorescence microscopy. In Figure 3 we show individual images (**Fig. 3A**) as well as averaged intensity distributions of end-tracking proteins on microtubules at two different Tea2 concentrations (20nM and 100nM). The averaged Tea2-Alexa647 intensity profiles demonstrated a specific shape: a shallow intensity increase starting at the microtubule seed, a constant average intensity along the microtubule lattice and a pronounced spike at the microtubule plus-end (**Fig. 3B**). Distributions of Tip1:GFP and Mal3-Alexa647 fluorescence along the microtubules followed the same overall profile as Tea2 (**Fig. 3C,D**). Interestingly, at the higher motor concentration Tip1 intensity increased more than Tea2 intensity itself both on the microtubule lattice and at microtubule ends (**Fig. 3B,C**). In contrast, the intensity of Mal3-Alexa647 did not change with Tea2 concentration (**Fig. 3D**).

**Figure 3:**
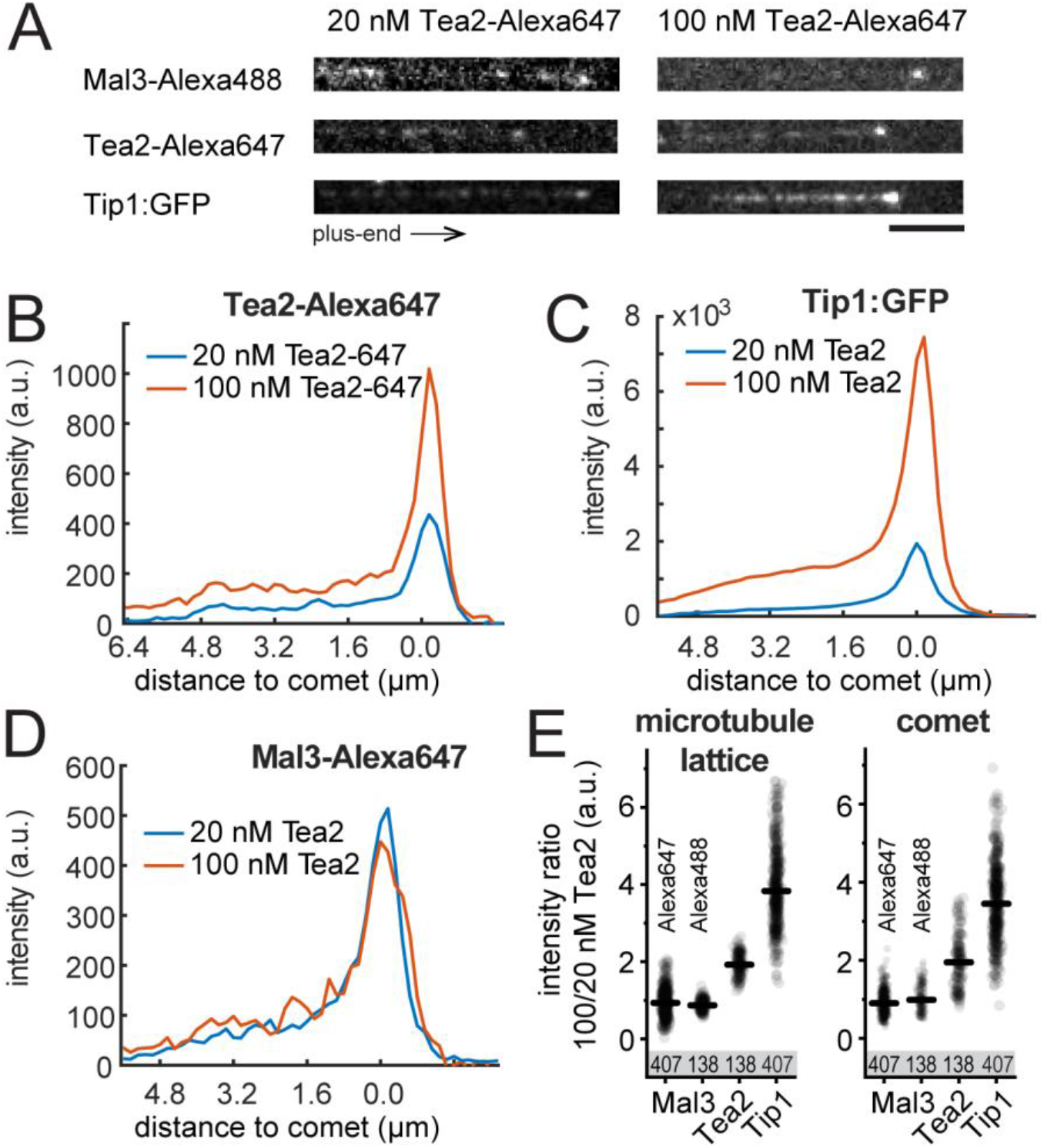
Non-stoichiometric accumulation of Tip1 at microtubule plus-ends. **(A)** Examples of individual Mal3, Tea2, and Tip1 intensity profiles at two different concentrations of Tea2 (20 nM and 100 nM).**(B)** Averaged Tea2-Alexa647 intensity profiles. **(C)** Averaged Tip1:GFP intensity profiles. **(D)** Averaged Mal3-Alexa647 intensity profiles. Data in (C) and (D) were extracted from the same experiment, whereas Tea2-Alexa647 data shown in (B) were recorded in a separate experiment which contained dark Tip1 and Mal3-Alexa488. **(E)** Ratio of lattice (left) and comet (right) intensities between 100 and 20 nM Tea2 for all data. The number of observed intensity profiles per condition is indicated.

We summarized the effect of Tea2 concentration on the end-accumulation of Mal3 and Tip1 by calculating the ratios of intensities between the two concentrations of Tea2 for Mal3 and Tip1 on the microtubule lattice and in the comet (**Fig. 3E**). An increase in Tea2 concentration had no influence on the amount of labelled Mal3 protein that localized at the microtubule plus end. On the other hand, Tip1:GFP localization to the plus end was disproportionately affected by Tea2 concentration. An increase from 20 nM to 100 nM Tea2 led to a roughly 4-fold increase of Tip1:GFP intensity at the plus end, whereas the Tea2 intensity itself was only increased by a factor of 2. Apparently, the amount of Tip1 that is present on the microtubule does not follow the density of motor proteins on the microtubule in a stoichiometric way. In fact, **Supplementary Fig. S3** shows that the presence of Tip1 responds in a non-linear way to the concentration of Tea2 over a wide range of concentrations (an effect that we observed previously but that is not understood (Fig. 1F in Taberner and Dogterom, 2019)). Note also that there is large variability in the Tip1 intensity between individual microtubules (**Supplementary Fig. S3A**) which we interpret as another sign that the accumulation of the cargo Tip1 is not limited by one-on-one interactions with motor proteins.

### Distinct domains of Mal3 contribute to formation of Mal3 comets and LLPS

Having established that our three-component network is capable of both droplet formation and non-stoichiometric protein accumulation at microtubule ends, we set out to elucidate the contributions of disordered protein regions to both comet formation and LLPS. As Mal3 is central to comet formation of all three proteins, we studied different truncations of Mal3. Fulllength Mal3 contains two folded domains: a calponin-homology (CH) domain, and an EB-homology domain (EB HD), and two intrinsically disordered regions: IDR1, which connects the CH domain to EB HD, and the C-terminal IDR2 (**Fig. 4A**). Note that the C-terminal IDR2 domain is not present in Mal3’s homolog EB1 which contains a much shorter negatively-charged C-terminal tail (Buey et al., 2011). We first focused on dissecting the contributions of these domains to formation of Mal3 comets on microtubule ends without Tea2 or Tip1, and in the absence of crowding conditions (**Fig. 4B,C**).

**Figure 4:**
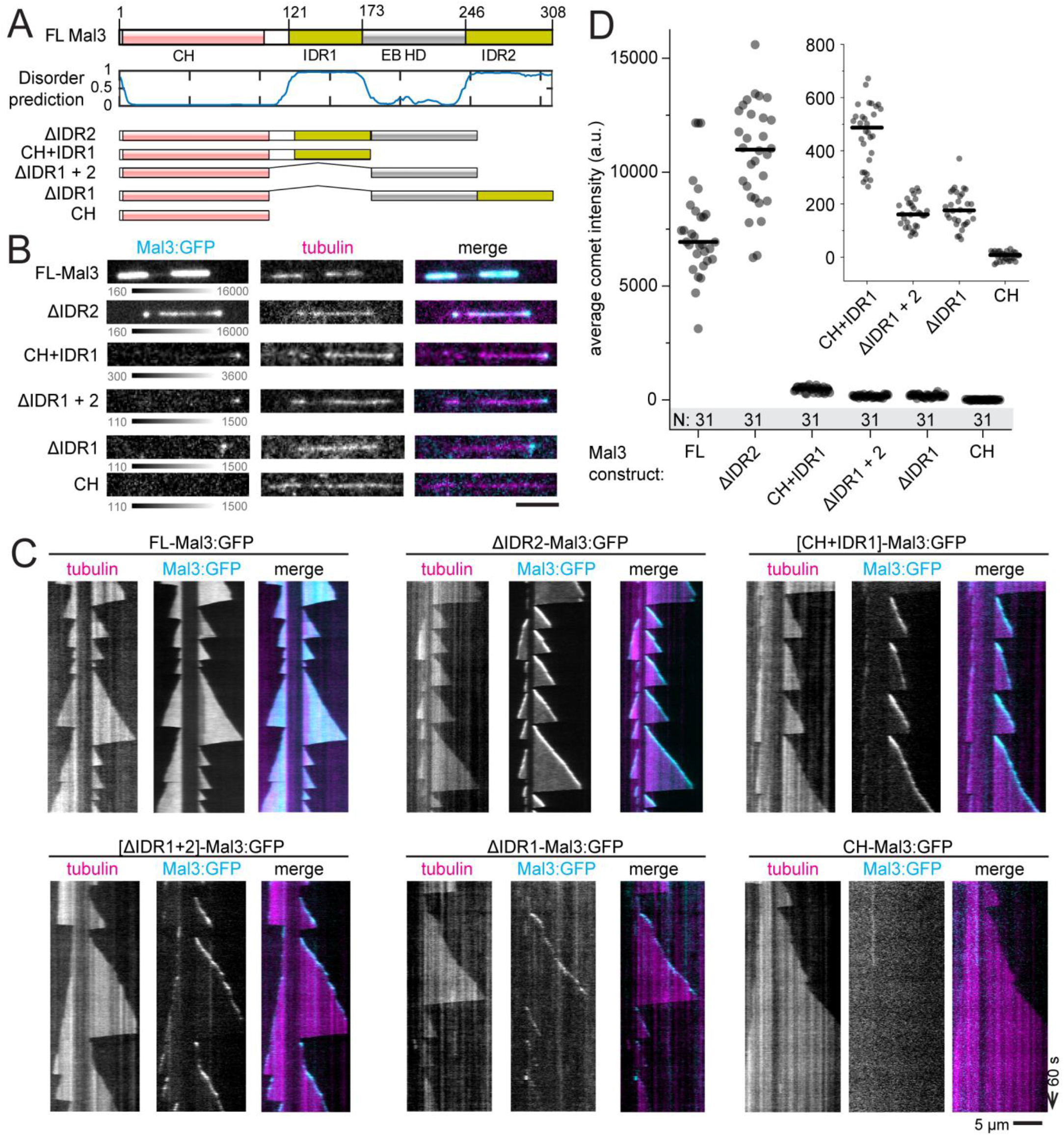
Mal3 IDR1 and EB HD are required for comet formation. **(A)** Pictorial representation of the full length Mal3 together with disorder prediction (Jones and Cozzetto, 2015) and the truncations used in this study. **(B)** 200 nM FL-Mal3 and Mal3 truncation mutants decorating the growing microtubule ends. **(C)** Kymographs for the end-tracking experiments presented in (B). **(D)** Average comet intensity for full length Mal3 and Mal3 truncates. Lines show median, numbers in the shaded area show N. Inset shows scaled up graph for four truncates with poorest binding. Scale bars: 5 µm and 60 sec.

In the absence of Tea2 and Tip1, 200 nM of full length Mal3:GFP coated the entire microtubule lattice without a clear saturation at the plus-end, in contrast to the full MPET network (compare **Fig. 4C** (top left) and **1B**, respectively). Mal3-ΔIDR2 showed a lower binding affinity to the microtubule lattice than full length Mal3 and formed slightly brighter comets at the microtubule ends (**Fig. 4C,D**). We did not observe any comet formation or lattice binding with Mal3 truncates containing only the CH domain (**Fig. 4B,C**). Other Mal3 truncates were binding mostly to the growing end, rather than the microtubule lattice. **Figure 4D** shows a comparison of the comet intensity observed with full length Mal3 and Mal3 truncations. All truncates except for Mal3-ΔIDR2 were binding very poorly to microtubule ends. We conclude that, in addition to previously described microtubule-binding through the CH-domain and the role of dimerization through the EB HD (Honnappa et al., 2005; Slep et al., 2005), IDR1 also contributes to efficient comet formation by Mal3. In contrast, IDR2 appears not to contribute to Mal3’s affinity to the microtubule end but only to its affinity to the lattice (potentially via Mal3 self-interactions; see below). Note that in previous work on EB1 truncates it was observed that removal of the C-terminal tail (where IDR2 is) led to stronger instead of weaker lattice binding (Buey et al., 2011). This effect was attributed to the removal of a short negatively charged section of the protein which is expected to destabilize electrostatic interactions with the positively charged microtubule lattice. While it is difficult to disentangle the effect of charge from the contribution of multivalent interactions, it should be noted that EB1 does not have a sizable, disordered region at its C-terminal end.

We next wondered which domains of Mal3 are important for the protein’s self-interactions under crowding conditions. When Mal3 truncates were incubated at a concentration 1 µM with 5% PEG 35k, we observed that domain deletions preventing comet formation on microtubules also prevented droplet formation (Fig. 5A). In addition, Mal3-ΔIDR2, which reduced microtubule lattice but not microtubule end-binding, also formed smaller condensates than the full length protein in the presence of a crowding agent. We conclude that IDR1 and EB HD are necessary both for Mal3 self-interactions and Mal3 interaction with the microtubule end, while IDR2 is also necessary for Mal3 self-interactions and interaction with the microtubule lattice, but not the microtubule end (**Fig. 4B,C** and **5A**).

**Figure 5:**
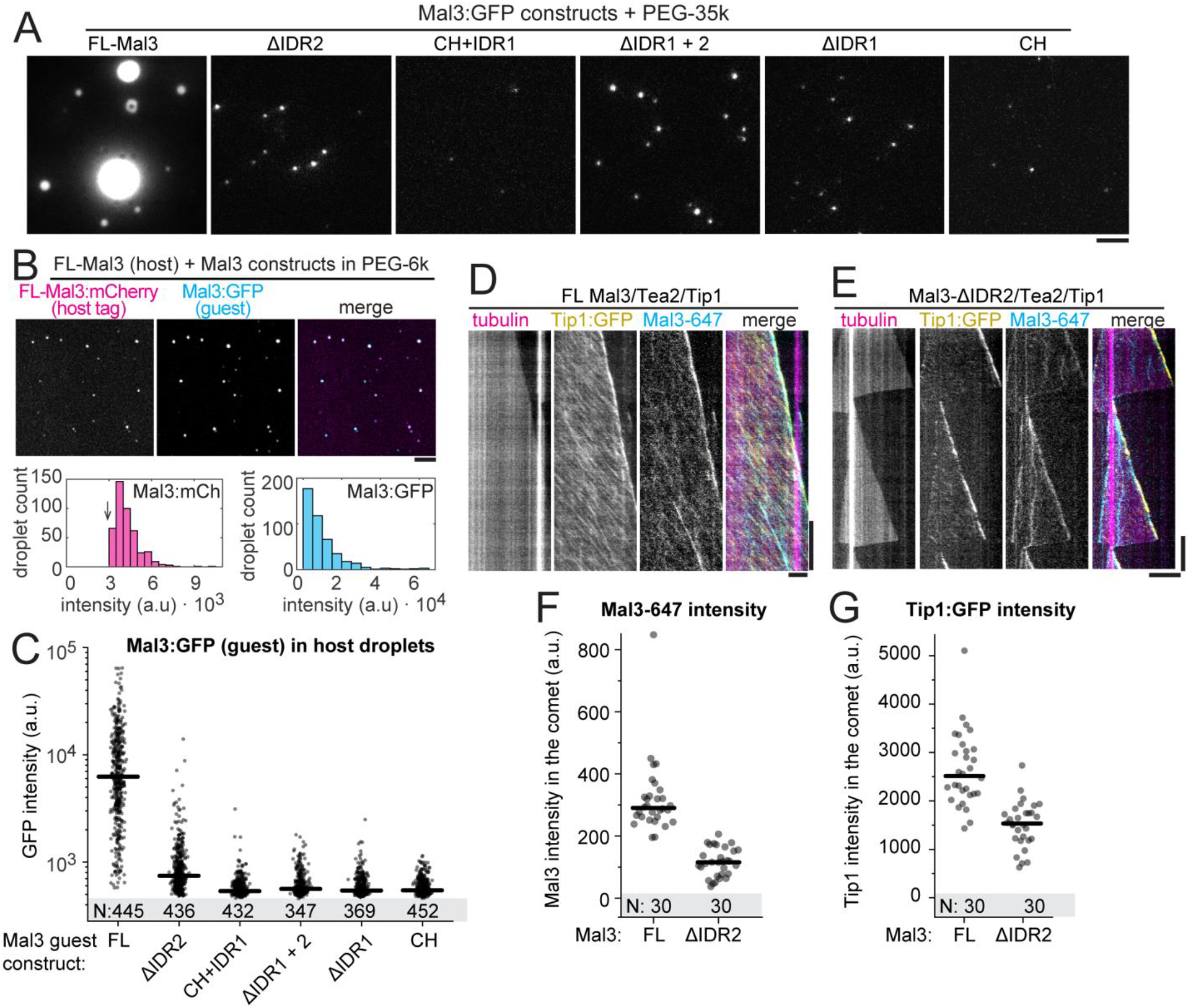
Distinct domains of Mal3 mediate comet formation and LLPS. **(A)** Condensates formed by full length Mal3 and Mal3 truncates (1 µM) in the presence of 5% PEG 35k. **(B)** Unlabelled FL-Mal3 (200 nM, host) tagged with FL-Mal3:mCherry (2 nM, tag) was allowed to recruit Mal3:GFP (2nM, guest) in presence of PEG-6k. Graphs show distributions of tag and host Mal3 intensities (arrow denotes the threshold applied for speckle detection in images). (**C**) Intensity of Mal3:GFP (full-length or truncated constructs) recruited to FL-Mal3 host droplets. Lines show median, numbers in the shaded area show N. **(D-E)** Kymographs showing end-tracking by Mal3/Tip1/Tea2 in presence of full-length Mal3 (D) or Mal3-ΔIDR2 (E) and in the absence of PEG. **(F-G)** Intensities of Mal3-Alexa647 (F) and Tip1:GFP (G) in the comets forming in presence of FL-Mal3 or Mal3-ΔIDR2, lines show median, numbers in the shaded area show N.

### The role of distinct Mal3 domains in the formation of multi-protein condensates and comets

To pin-point the interactions between Mal3, Tea2, and Tip1 under crowding conditions we next designed a guest-host assay (**Fig. 5B,C** and **Supplementary Fig. S4**). Condensates consisted of either of the three full length proteins as hosts (Mal3, Tea2-Alexa647, or Tip1), and were formed by incubation with 10% PEG 6k, and 2 nM Mal3:GFP construct as the guest. When non-fluorescent Mal3 and Tip1 were used, we additionally added 2 nM full length Mal3:mCherry as a tag to visualize the host condensates independent of Mal3:GFP construct localization. Fig. 5B shows the outcome of a typical experiment, where in this case non-fluorescent Mal3 was the host (tagged with Mal3:mCherry), and full length Mal3:GFP was added as guest. As can be seen in **Fig. 5C**, deletion of any disordered region from Mal3:GFP prevented its recruitment to the Mal3 host condensate, reinforcing our conclusion that both IDR1 and IDR2 are important for Mal3-Mal3 interactions in crowding conditions.

We observed a direct interaction between full length Tea2 and Mal3 in crowding conditions in the absence of Tip1 (**Supplementary Fig. S4A**). However, Mal3 constructs lacking IDR1 or IDR2 were recruited poorly to Tea2-Alexa647 host condensates (**Supplementary Fig. S4A**). Deletion of EB HD further disrupted recruitment of Mal3 to the Tea2 host condensates. These data indicate that crowding conditions strengthen Tea2-Mal3 interactions and that these interactions rely on the disordered regions in Mal3 as well as the EB HD. Finally we used unlabeled Tip1 as the host condensate (**Supplementary Fig. S4B**) and Mal3:GFP truncates as the guests. We again observed that Tip1 condensates predominantly recruited full-length Mal3:GFP and, to a much lesser extent Mal3-ΔIDR2, but failed to recruit the Mal3 constructs lacking the EB homology domain or IDR1.

We finally set out to correlate the recruitment behavior observed in the guest-host assays with the capacity of truncated Mal3 constructs to couple Tip1/Tea2 transport to plus-end tracking on dynamic microtubules. Using Tip1:GFP fluorescence as a readout, we observed that Mal3 constructs lacking either IDR1, EB homology domain, or their combinations, failed to recruit Tip1:GFP to microtubules altogether (**Supplementary Fig. S5**). Mal3-ΔIDR2 was still able to support Tip1 localization at microtubule ends, but unlike full length Mal3, it did not colocalize with Tea2/Tip1 transported along the microtubule lattice (**Fig. 5D,E**). Furthermore, the intensity of both Mal3-ΔIDR2 and Tip1 in the comets was reduced compared with full length Mal3 (**Fig. 5F,G**).

Together, the analysis of Mal3 truncations leads us to conclude that robust three-component comets are formed by a combination of different molecular mechanisms. Mal3 interaction with itself, Mal3 interaction with the microtubule lattice, as well as Mal3 co-localization with motor tracks requires each of Mal3’s IDRs. The formation of Mal3/Tip1 comets requires only Mal3 IDR1 and EB HD, but the additional presence of IDR2 enhances the motor-dependent accumulation of Tip1 at growing microtubule ends. It thus appears that Mal3 self-interactions are needed to promote non-stoichiometric Tea2/Tip1 transport on the microtubule lattice.

### Theoretical models for comet formation

To help understand the relative contributions of protein self-interactions and motor activity to efficient comet formation, we next turned to stochastic simulations. It should be noted that these simulations are not designed to exactly reproduce the experimental situation which is quite complex. The simulations specifically focus on the ability of motor proteins (Tea2) to enhance the presence of cargo molecules (Tip1) at microtubule ends in different scenarios (with/without end-specific motor behavior and with/without the ability of Tip1 to form clusters). They for example do not take into account the fact Tip1 may also accumulate at microtubule ends via transport-independent direct recruitment by the autonomous end-tracker Mal3 (see e.g. **Fig. 5E**, but also Fig. 1C in Taberner and Dogterom, 2019).

We simulated a set of different models where the microtubule plus end is represented by a growing one-dimensional lattice (Melbinger et al., 2012; Reese et al., 2014) and each lattice site corresponds to one tubulin heterodimer. The molecular motor Tea2 is represented by particles (circles in **Fig. 6A**) which bind to and unbind from the lattice and hop towards the plus end (Leduc et al., 2012; Lipowsky et al., 2001; Parmeggiani et al., 2003), where each lattice site can be occupied by only one motor. The molecular cargo is represented by a second set of particles that can bind to and unbind from the motor particles (squares in **Fig. 6A**) but cannot bind to the microtubule lattice. We consider the effects of Mal3 indirectly by assuming different motor/cargo behaviors at microtubule end and lattice and by studying different scenarios for cargo oligomerization. We do not treat Mal3 as separate particle species, because experiments showed that Mal3 localization is not significantly affected by motor concentration (**Fig. 3D** and **Supplementary Fig. S3B**).

**Figure 6:**
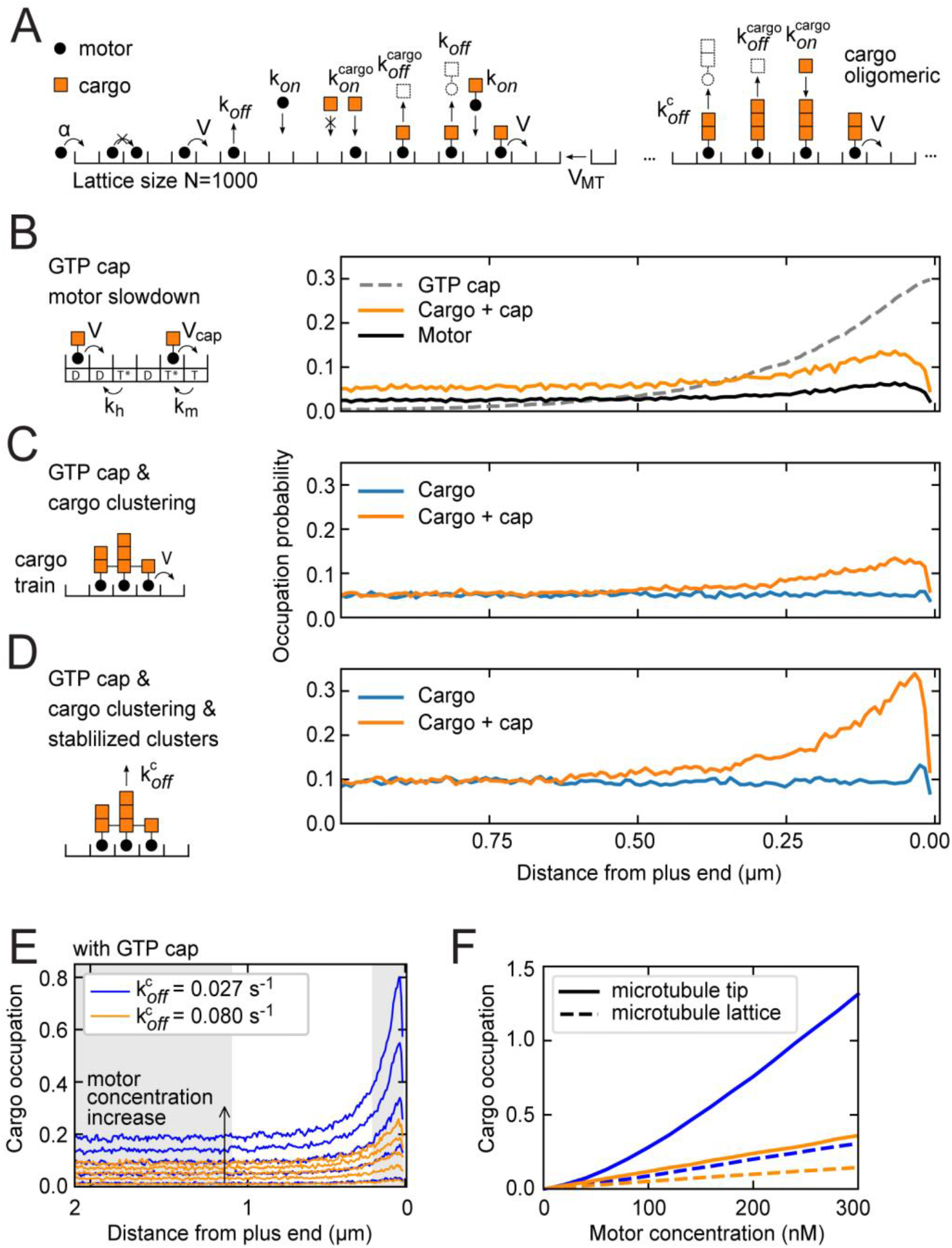
Theoretical models and stochastic simulations. **(A)** Cartoon of a lattice model for motor transport with cargo oligomerization. Reactions include motor movement, attachment, detachment, a growing lattice, the binding and unbinding of cargo to bound motors as well as a cargo multi-layer absorption/desorption process. Model parameters are provided in the Supplementary Table S2. **(B)** Nucleotide dependent motor slowdown was implemented by including a GTP cap and two-step hydrolysis. The GTP profile decays exponentially (dashed line, normalized to 0.3) and the amounts of motors and cargos at the lattice end increase.**(B)** Interactions between cargo particles was implemented such that neighboring cargo particles form a cargo train which induces coherent movement of cargo-motor clusters. Cargo clustering does not lead to increasing amounts of cargo at the lattice end unless a GTP cap is implemented as in (B). **(D)** The stability of cargo clusters was increased by dynamically enhancing the motor dwell time in cargo clusters. This mechanism alone did not result in significant accumulation of cargo, similar to panel (C). However, cargo accumulation increased sharply in combination with a GTP cap. Motor concentration in (B-D) corresponds to ∼100 nM Tea2 in experiments. **(E)** Density profiles of stabilized cargo clusters (blue lines) and independent cargo clusters (orange lines) for a range of motor concentrations between 20 and 180 nM. **(F)** The average cargo occupation is shown depending on motor concentration for the microtubule lattice (shaded area in panel (E) on the left; dashed lines) and the microtubule tip (shaded area in panel (E) on the right corresponding to ∼200 nm; solid lines).

The microtubule-motor-cargo kinetics described above are the starting point of our modeling approach. The phenomenology of this model is well known for motor transport in the absence of cargo (Parmeggiani et al., 2004; Reese et al., 2011), and because the binding/unbinding of cargo is an equilibrium process, it is not expected that simple one-on-one cargo binding changes any characteristic of the model. Here we are however interested in the situation where multiple cargo molecules can bind to each other, allowing cargo-oligomerization on top of single motors (**Fig. 6A**) as well as lateral cluster formation between cargo associated with different molecular motors (**Fig. 6C,D**) (Brunauer and Emmett, 1938; Mitchison, 2020). This is inspired by our experimental observations in guest-host and end-tracking assays, but also supported by other evidence that Tip1 may be able to oligomerize (Chen et al., 2019; Taberner and Dogterom, 2019) and structural data which suggests head-tail interactions between Tip1’s CAP-Gly domain and its C-terminal zinc finger domain (Hayashi et al., 2007; Steinmetz and Akhmanova, 2008). We chose quantitative model parameters where possible (see Materials and Methods and Supplementary Table S2) which also reflect the experimental observation that motors fall off the microtubule end at approximately their stepping rate in the absence of Mal3 (**Supplementary Fig. S6F**) (Bieling et al., 2007b; Munteanu, 2008). We then set out to find conditions leading to increased cargo accumulation at microtubule ends as a result of motor transport, end-dependent motor-microtubule interactions due to end-specific microtubule properties (e.g. the nucleotide state), and/or cargo clustering.

First, we investigated the effect of the nucleotide state at the microtubule plus end on motor dynamics (in the absence of lateral cargo interactions). In experiments we observed that when we mimicked the hydrolysis transition state of GTP using GTPγS microtubules, motor transport effectively slowed down by a factor of two (**Supplementary Fig. S6A-C**). This observation gives rise to the hypothesis that cargo accumulation could be the result of motor slowdown at the plus end (**Fig. 6B**). Indeed, simulations show that slower movement of motors at the microtubule end leads to an accumulation of motors and cargo at the plus end (**Fig. 6B**). The effect arises from a backlog, or traffic jam at the transition from the fast to the slow parts of the lattice (indicated by the dashed line **Fig. 6B**).

The second scenario we investigated considered the additional lateral interaction between neighboring motor-bound cargo molecules (**Fig. 6C**). To this end we implemented clustering of cargo particles into oligomeric cargo trains, which moved synchronously in the simulation (Bunzarova et al., 2019). Cargo trains increased the effective flux of cargo on the microtubule (due to synchronous stepping of clusters) but did not lead to accumulation of cargo at the microtubule end under the given conditions (**Fig. 6C**). An accumulation of cargo is recovered by introducing motor slowdown at the microtubule end as in **Fig. 6B**.

As a last plausible modification to the model, we explored the effect of increasing the stability of cargo clusters. In our model the stability of cargo clusters is determined by the detachment rate of a motor (k_off_ = 0.08 s^-1^ with bound cargo), and by the detachment of cargos (k_off_^cargo^ = 0.01 s^-1^). To assess the importance of cargo cluster stability we made the dwell time of motors dependent on the presence of neighboring motors with cargo. The mechanism was implemented such that motors with cargo and at least one neighboring motor with cargo, remained bound to the lattice at a dwell time that was longer as compared to motors which were not part of a cargo cluster (**Fig. 6D**). Even a threefold increase in dwell time was not sufficient to cause accumulation of cargo at the microtubule end (**Fig. 6D**), similar to our previous observation (**Fig. 6C**). However, if in addition the end-dependent slowing down was implemented, the model resulted in pronounced cargo accumulation at the microtubule end (**Fig. 6D**). Thus, a mechanism in which motor dwell time is dynamically increased through lateral cargo clustering on the microtubule lattice appears to be, in combination with motor slow down at microtubule ends, a very efficient way to accumulate cargo at the microtubule end. Note that such a stabilization of oligomeric clusters also increases the lattice occupancy away from the microtubule end (cf. **Fig. 6D,E**). The computational model also allows us to study the effect of cargo clustering on cargo transfer events as experimentally observed under crowding conditions (**Fig. 1H**). To this end we simulated cargo transfer events with different cluster-dependent motor dwell times as introduced above (**Fig. 6D**). For increasing motor dwell times within cargo clusters, clusters travel increasingly long distances along the lattice (**Supplementary Fig. S6G**,**D**).

Finally, to help understand the motor concentration-dependent increase in accumulation of cargo at microtubule plus-ends in our experiments (**Fig. 3E** and **Supplementary Fig. S3**), we investigated how the accumulation of the cargo depends on the concentration of motors in the model. To this end we simulated a range of motor concentrations from 20 to 180 nM (**Fig. 6E** and **Supplementary Fig. S6E**) for the scenarios shown in **Fig. 6C&D**. Only the scenario in which motors slow down at the microtubule end, *and* cargo clusters are stabilized by lateral interactions resulted in a non-linear accumulation of cargo at the microtubule end (**Fig. 6E,F**).

Taken together, our models show that the combination of motor slow-down at microtubule ends, lateral interactions between neighboring motor-cargo clusters, and increased motor processivity in cargo clusters may lead to robust formation of comets. Notably, in such a scenario the comet size depends non-linearly on the motor concentration providing an effective end-localization mechanism for motor driven cargo on microtubules.

## Discussion

In this study we systematically dissected the role of multivalent interactions within the microtubule plus-end tracking network reconstituted *in vitro* using recombinant Mal3, Tip1, and Tea2 from *S. pombe*. We found that *in vitro* molecular crowding agents, such as PEG, drove these proteins into spherical droplets that displayed liquid-like properties: they fused with each other over time, wetted the microtubule surface, transferred from one microtubule to another and were transported by Tea2 motor activity towards microtubule plus-ends. This behaviour shows similarity to the previously observed transfer of end-tracking protein clusters from a microtubule end to a solid barrier (Taberner and Dogterom, 2019), and might be relevant *in vivo* for the cortical deposition of polarity markers that are crucial for the physiology of fission yeast such as Tea1, Tea4, and Tea3 in addition to Mal3, Tea2, and Tip1 (Behrens and Nurse, 2002; Feierbach et al., 2004; Meadows et al., 2018; Snaith et al., 2005; Snaith and Sawin, 2003).

We found that under crowding conditions Mal3, Tip1, and Tea2 coexisted in the same condensed phase, confirming previous reports of Mal3-Tea2, Mal3-Tip1 and Tip1-Tea2 interactions in the absence of microtubules (Browning and Hackney, 2005; Busch et al., 2004; Busch and Brunner, 2004). However, previously it remained unclear which domains of Mal3 mediate these interactions. Here we show that IDR1 and IDR2 in Mal3 are responsible for multivalent interactions with Tip1 and Tea2 in the absence of microtubules: Mal3-ΔIDR1 and Mal3-ΔIDR2 were recruited poorly to condensates made of full length Mal3, Tea2, or Tip1 (Fig. 5C, Supplementary Fig. S4A,B). Deletion of the same disordered regions of Mal3 impaired formation of Mal3 droplets in crowding conditions (**Fig. 5A**), in accordance with the idea that disordered regions are the main drivers of LLPS (Alberti et al., 2019; Wang et al., 2018). We further found that Mal3 IDR1, in combination with EB HD, is crucial for Mal3’s accumulation at growing microtubule plus-ends. Deletion of IDR1 leads to 1) reduced intensity of Mal3 end-tracking comets, and 2) absence of Tip1 accumulation at the microtubule plus ends (**Fig. 4, Supplementary Fig. S5**).

Importantly, Mal3-ΔIDR2, for which we also observed severely impaired droplet formation and interaction with Tip1 and Tea2 in crowding conditions, was nevertheless able to form comets at growing microtubule ends (**Fig. 4B-D**) and recruit Tip1 to these comets (**Fig. 5E,G**). However, both Mal3 and Tip1 intensity were reduced in comets formed by Mal3-ΔIDR2, and Mal3-ΔIDR2 failed to associate with Tea2 transport (**Fig. 5E,G**) in addition to showing reduced lattice binding (**Fig. 4B,C**). We conclude that robust motor-driven transport and accumulation of Tip1 at microtubule ends depends on both Mal3 IDR1 and IDR2, leading to the suggestion that Mal3 self-interactions responsible for LLPS are also responsible for protein interactions in the network-like structures observed at microtubule ends in cryo-ET (**Fig. 2E-F**).

The question that then remains is whether the network-like structures observed in cryo-ET in the absence of crowding agents show characteristics of liquid-like droplets and/or whether this is expected to be the case for end-tracking complexes *in vivo*. Clearly, PEG-driven droplets in the absence of microtubules displayed a characteristic dense internal grain in cryo-ET (**Fig. 2B,C**) that was not seen in the microtubule end-tracking comets. The comets appeared as much more loosely structured densities (**Fig. 2E,F**) which did not extend further than 55 nm from the microtubule lattice (28 nm on average, **Fig. 2G,H**). Given the estimated dimensions of Mal3 (3×6×10 nm) (Matsuo et al., 2016; von Loeffelholz et al., 2017), Tea2 (4×4×7 nm) and Tip1 (predicted 40nm-long coiled coil), it is therefore technically possible that all the molecules inside the comet are directly interacting with microtubule surface. On the other hand, it is also possible that the loose network represents a liquid-like structure where multiple dynamic, weak interactions between its components facilitate the observed non-stoichiometric accumulation of plus-end trackers and allow the network to behave as a protein cluster (Taberner and Dogterom, 2019). *In vivo*, where crowding effects are stronger than in the conditions of our cryo-ET experiments, these clusters may again appear as dense droplets as we observed in the presence of PEG. For completeness, it should furthermore be noted that preserving dense droplets at the ends of microtubules during our cryo-ET sample preparation may be technically challenging, potentially limiting our ability to properly visualize these structures.

In conclusion, the suggestion that arises from our study is that microtubule ends may act as platforms where multivalent interactions act to condense large numbers of microtubule-associated proteins into functional complexes. In parallel to our work on motor-dependent accumulation of fission yeast proteins, observations in three other biological systems are pointing in this same direction. Meier et al. report on the formation and biological relevance of Kar9 nano-droplets at the ends of specialized microtubules in budding yeast, Miesch et al. describe the formation of human EB3 and Clip170 droplets at microtubule ends leading to enhanced local tubulin concentration, and Song et al. describe the importance of EB1 condensation for chromosome segregation during mitosis. The notion of liquid-liquid phase separation at microtubule ends is thus emerging as a general organizing principle that may explain how different end-tracking proteins may (simultaneously) associate with microtubule ends and perform their wide range of biological functions.

## Materials and Methods

### Protein expression, purification, and labelling

Full length *Schizosaccharomyces pombe* Mal3 and all of its derivatives (*i*.*e*. truncates and sfGFP fusions) were expressed with an N-terminal His8 tag followed by a 3C protease site from a pBAD-TOPO derived plasmid in *Escherichia coli* ER2566 cells (New England Biolabs, *fhuA2 lacZ::T7 gene1 [lon] ompT gal sulA11 R(mcr73::miniTn10--Tet*^*S*^*)2 [dcm] R(zgb-210::Tn10--Tet*^*S*^*) endA1 Δ(mcrCmrr)114::IS10)*. Mal3(-truncates) were covalently linked to superfolder-GFP by a flexible ASTGILGAPSGGGATAGAGGAGGPAGLINPGGSTSSRAAEIWPAS “happy linker” sequence. Cells were grown at 37 °C in baffled flasks on LB supplemented with 100 µg/ml ampicillin, expression was induced at an OD_600_ of 0.6, and cells were harvested after 3 hours (8 min 4500 rpm, JLA8.1000 rotor). After washing the cells in PBS they were lysed using a French Press (Constant Systems) at 20 kpsi, 4 °C and unbroken cells, debris and aggregates were pelleted in a Ti45 rotor (30 min, 40.000 rpm, 4 °C). The lysate was applied to 2 ml Talon Superflow resin (Clontech) pre-equilibrated with buffer A (20 mM Tris/HCl pH 7.5, 200 mM NaCl, 5% (w/v) glycerol), and incubated for one hour while rotating at 4 °C. Subsequently, the resin was washed with 50 ml of buffer A supplemented with 0.1% Tween20, 50 ml of buffer A supplemented with additional 500 mM NaCl, and finally Mal3 was eluted in 10 ml of buffer A supplemented with 1 mM β-mercaptoethanol and homemade 3C protease. Proteins were concentrated using a Vivaspin centrifugal concentrator (10 kDa cut-off) and further purified by size exclusion chromatography (SEC) on a Superdex 200 Increase 10/300 column pre-equilibrated with buffer B (20 mM Tris/HCl, 100 mM NaCl, 5% (w/v) glycerol).

Mal3 was labelled by dialyzing ∼1 mg of protein into buffer C (80 mM PIPES pH 6.8, 1 mM MgCl_2_ 1 mM EGTA, 100 mM NaCl) and incubating for one hour at room temperature with 140 µM Alexa Fluor 488 TFP ester or Alexa Fluor 647 TFP ester (Thermo Fisher). After quenching the reaction with excess Tris/HCl the free label was removed by SEC on a Superdex 200 Increase 10/300 column pre-equilibrated with buffer B.

Full length *Schizosaccharomyces pombe* Tea2 was expressed with an N-terminal Z-tag followed by a TEV protease recognition site, and purified essentially as described (Bieling et al., 2007a), but with the following modifications: after washing of the Talon resin with 15 mM imidazole in buffer D (50 mM KPi pH 8.0, 400 mM NaCl, 2 mM MgCl_2_, 0.2 mM MgATP, 0.05 mM TCEP), Tea2 was eluted in buffer D supplemented with homemade 3C protease by taking advantage of cross-reactivity with the TEV recognition site. Following concentration using a Vivaspin centrifugal concentrator (10 kDa cut-off), Tea2 was labelled with 138 uM Alexa Fluor 647 NHS ester (Thermo Fisher) by incubating 30 minutes at room temperature. After quenching the reaction with excess Tris/HCl the free label was removed by SEC on a Superdex 200 Increase 10/300 column preequilibrated with buffer D. Unlabelled Tea2 was applied to the SEC column directly after concentrating.

### Flow cell preparation

Cover slips and glass slides were cleaned using base Piranha (NH_4_OH:H_2_O_2_ in 3:1 at 75°C) for 10 min and sonicated in milliQ water for 5 min. Flow cells were prepared by sandwiching two strips of parafilms between the glass slid and the coverslip. The strips were placed about 3-5 mm apart approximately from each other. The flow cell was then placed on top of a hot plate, kept at 120 °C, to let the parafilm melt and seal the glass slid with the coverslip.

### Microtubule Biochemistry

#### a) GMPCPP-stabilized microtubule seeds

Microtubule seeds were prepared by two cycles of polymerization with GMPCPP in MRB80 buffer (80 mM Pipes, 4 mM MgCl2, 1 mM ethylene glycol tetraacetic acid (EGTA), pH 6.8). First 20 µM tubulin (25% HyLite 647, 10% Biotinalyted and 65% unlabelled) was polymerised in the presence of 1 mM GMPCPP (NU-405 Jena BioScience, Jena, Germany) at 37 °C for 30 min. The mix was centrifuged for 5 min at 30 psi with an air-driven ultracentrifuge, airfuge (Beckman Coulter Inc., Brea, California, USA) and the pellet was resuspended in MRB80 (80% of the initial volume) and kept on ice for 20 min for depolymerisation. For the second polymerization step, again 1 mM of GMPCPP was added to the mix and the mix was incubated at 37 °C for another 30 min. After 30 minutes of incubation the mix was ultra-centrifuged using airfuge and the pellet was resuspended in 50 µl MRB80 with 10% glycerol. The seeds thus prepared were aliqoted, flash frozen in liquid nitrogen and stored at − 80°C.

#### (b) End-tracking reconstitution assays

To functionalize the glass surface, the channels in the flow cells were first filled with 0.2 mg/ml PLL(20)-g[3.1]-PEG(2)/-PEG(3.4)-biotin(17.5%) (SUSOS AG) then 0.1 mg/ml neutravidin and finally with k-casein (Sigma). 10 min incubation at room temperature was maintained before the subsequent steps. The channels were then washed with MRB80 and incubated with biotinylated seeds for 5 min. After 5 min the reaction mix was added to the channels. The channels were sealed with VALAP before starting the observations on the microscope to avoid evaporation.

To reconstitute plus-end-tracking assays with full length Mal3 and Mal3 truncates, the reaction n mix contained 200 nM Mal3/Mal3 truncate, 20 nM Tea2 and 150 nM Tip1 in MRB80 buffer containing 14.5 µM tubulin, 0.5 µM rhodamine tubulin, 50 mM KCl, 0.5 mg/ml k-casein, 0.4 mg/ml glucose oxidase, 50 mM catalase, 0.1% methyl-cellulose, 1 mM GTP, and 2 mM ATP.

### Phase separation assays

#### a) Condensates on dynamic microtubules

The assay was performed in two steps. In the first step a dynamic microtubule assay was set up in a flow cell and in the second step condensates were added. In order to set up a dynamic microtubule assay, a reaction mix with 14.5 µM tubulin, 0.5 µM rhodamine tubulin, 50 mM KCl, 0.5 mg/ml k-casein, 0.4 mg/ml glucose oxidase, 50 mM catalase, 0.1% methylcellulose, 1 mM GTP, and 2 mM ATP in MRB80. The flow cell was then left for incubation at 37 °C for 15 min. After 15 min the tubulin was washed off using MRB80 (per-warmed at 37 °C) and condensates were added to the flow cell immediately to the flow cell. The condensates were prepared by incubating 200 nM Mal3, 20 nM Tea2 and 150 nM Tip1 in MRB80 buffer containing 50 mM KCl, 0.5 mg/ml k-casein, 0.4 mg/ml glucose oxidase, 50 mM catalase, 0.1% methylcellulose, 1 mM GTP, and 2 mM ATP on ice for 1 hr with 5% PEG 35k.

#### b) Guest-Host experiments

Cover slips were cleaned as described above. Glass slides were cleaned in a 250 ml beaker with a custom-made Teflon rack by repeated (2x) sonication and washing steps as follows 1% Hellmanex (10 minutes), MilliQ Water (5 minutes), 70% Ethanol (10 minutes), MiliQ (5 minutes), and stored in the beaker with MiliQ covered by parafilm. Before use, slides were rinsed with MiliQ and dried with N_2_.

Flow cells were prepared by cutting six channels into a piece of parafilm with a razor blade. The parafilm was sandwich between the clean glass slide and the cover glass and heated on a piece of aluminum foil on top of a 120 °C hot plate until the parafilm melted and cover glass was gently pressed with tweezers to assure that channels were sealed off well. The parafilm overhangs were removed with the blade while the glass was still hot. After cooling to room temperature the channels were incubated for 10 minutes with 0.2 mg/ml PLL(20)-g[3.1]-PEG(2) (SUSO AG), rinsed, and incubated for 10 minutes with 0.5 mg/ml k-casein (Sigma), all solutions were MTB80 buffer.

The guest-host condensates were prepared on ice by first eluting all proteins into MRB80 buffer containing 250 mM KCl, and then further diluting them into the reaction mixture, at a final composition of 1x MRB80, 50 mM KCl, 10% PEG6k, and freshly thawed 2 mM ATP, 1 mM GTP, 2 mM DTT, and 0.5 mM β-mercaptoethanol. Solutions were clarified for 5 min using airfuge and kept on ice for 15 more minutes before being transferred into flow cells. Imaging occurred approximately 30 minutes after mixing. Mal3 and Tip1 host condensates were prepared with 200 nM FL Mal3 or 215 nM Tip1, 2 nM FL Mal3:mCherry and 2 nM of each of the constructs (Fig. 3a). Tea2 host condensates were prepare from 200 mM Tea2-Alexa647 and 2 nM of each of the constructs. Experiments for each host protein were conducted in parallel. Image acquisition was performed using spinning disc confocal microscopy (CSU-W1, Yokogawa; Ilas2, Roper Scientific) with the scanning slide module in the Ilas2 software.

### Cryo-electron tomography

To study PEG-driven droplets, a solution containing 200 nM Mal3, 150 nM Tip1 and 80 nM Tea2 was incubated with 10% of PEG-8k in MRB80; 4 µL of this solution was mixed with 5 nm gold nanoparticles (OD50, final dilution 1:20) and added to freshly glow-discharged copper grids with R2/2 Quantifoil film. The grid was blotted from the back side for 4-6s in a Leica EM GP plunger and immediately plunge-frozen in liquid ethane.

To reconstitute comet formation, we used copper mesh grids with holey SiO film (SPI Supplies), coated with 5 nm gold on one side. The grids were treated with oxygen plasma for 2 min and immediately submerged in Plus-One Repel Silane solution (GE Life Sciences) for 3 min, then washed in ethanol and dried. A silanized grid was incubated in a drop of anti-DIG IgG (0.2µM, Roche), washed with MRB80, incubated in a drop of 1% Pluronic F-127 and washed again with MRB80. The passivated grid was then taken into the chamber of the Leica EM GP2 plunger equilibrated at 95% relative humidity and 26°C. Inside the chamber, GMPCPP-stabilized, DIG-labelled microtubule seeds were added to the grid for 1 min followed by a wash with MRB80 supplemented by 0.5 mg/ml κ-casein, and finally a 4 µL drop of a solution containing 200 nM Mal3, 150 nM Tip1 and 80 nM Tea2 in MRB80 supplemented with 25 µM tubulin, 0.01% tween-20, 2 mM ATP, 1 mM GTP and 1 mM DTT. The microtubules were allowed to grow for 7 min, after which 5 nm gold nanoparticles were added (OD50, final dilution 1:20), the grid was blotted from the back side for 3-4 s and immediately plunge-frozen in liquid ethane. All grids were stored in closed boxes in liquid nitrogen until further use.

Tilt-series were recorded on a JEM3200FSC microscope (JEOL) equipped with a K2 Summit direct electron detector (Gatan) and an in-column energy filter operated in zero-loss imaging mode with a 30 eV slit width. Images were recorded at 300 kV with a nominal magnification of 10000x, resulting in the pixel size of 3.668 Å at the specimen level. Automated image acquisition was performed using SerialEM software (Mastronarde, 2005) with a custom-written script, recording bidirectional tilt series ranging from 0° to ±60° with tilt increment of 2°; a total dose of 100 e^−^Å^2^ and the target defocus set to −4 µm. Individual frames were aligned using MotionCor2 (Zheng et al., 2017), and then split into odd and even frame stacks. Tilt series alignment and tomographic reconstructions were performed with the IMOD software package using gold beads as fiducial markers (Kremer et al., 1996). Final tomographic volumes were binned twofold and subsequently denoised using the cryoCARE procedure (Buchholz et al., 2019). For this, 3D reconstruction was performed on aligned sets of odd and even frame stacks with identical IMOD parameters. The full even and odd tomograms obtained in this way were then split into sub-volumes for network training, and eventually full volumes were denoised. The images presented in Fig. 5 were obtained from a voxel-wise average of odd and even denoised tomograms. Automated segmentation of binned and denoised cryo-tomograms was performed using the *tomoseg* module of EMAN2 v.2.2 (Chen et al., 2017) and visualized using UCSF Chimera (Pettersen et al., 2004).

### TIRF Microscopy

Imaging was performed using an inverted Nikon Eclipse Ti-E microscope with perfect focus system, an oil immersion objective (Nikon Plan Apo λ 100× NA 1.45), using two EMCCD cameras (Photometrics Evolve 512), which are mounted on a spinning disc unit (CSU-W1, Yokogawa). TIRF illumination was generated with the FRAP/TIRF system Ilas2 (Roper Scientific). A custom-made objective heater was used for temperature control of the samples. The imaging software used was Metamorph 7.8.8.0 with system specific routines (Ilas2) for streaming, time lapse, and scanning slide acquisition.

### Stochastic simulations

Stochastic simulations were performed using Gillespie’s algorithm (Gillespie, 1976) on the TU Delft Applied Science in-house linux cluster using an implementation in C++. The eight different models were simulated, tested, and prepared independently. The data presented shows the closest quantitative agreement with experimental conditions (see Supplementary Table S2 for a comparison) and corresponds a low density regime (LD phase) in terms of the TASEP/LK model on growing microtubules (Melbinger et al., 2012).

System size was *N* = 1000 lattice sites, each corresponding to the size of one tubulin heterodimer (8.4 nm). Simulations were equilibrated for 10^5^ sec before 10^4^ motor and cargo distributions were recorded in time intervals corresponding to the time it takes for one motor to traverse the system (∼50 sec). Equilibration times were particularly critical for cargo clustering conditions, since the motor distributions generically deviate from their classical equilibrium due to the aggregation and fragmentation kinetics, as seen in similar systems (Bunzarova et al., 2019). Data analysis and plotting was performed using custom programs and scripts written in C++ and Python (Matplotlib). Details regarding all model parameters and corresponding experimental values can be found in Supplementary Table S2 in the Supporting Material.

### Data Analysis

#### a) Preparation of density profiles

Kymographs were extracted from background subtracted TIRF microscopy data in a semi-automated way using ImageJ (50 pixel rolling ball radius). Image projections were used to identify dynamic microtubules in movies (function Z Project with option standard deviation) and positional data was stored in the form of linear regions (thickness 9 pixel) using the ImageJ ROI manager. Saved ROIs were used to automatically generate kymographs.

Subsequently a Matlab (R2018b) script was used to analyze kymographs (dual color where necessary) in a semi-automated way. The script allows to manually mark regions of growing microtubules with comets as polygons (typically triangular), generates a mask thereof, and extracts the corresponding intensity profiles from the underlying images. The intensity profiles are saved per experimental condition for further processing.

In a separate step the intensity profiles were sorted into sets by length using a binning of (± 0.64 µm). The set of profiles was aligned by finding alignments which minimize the standard deviation of the sum of differences between a randomly chosen first intensity profile and every other profile in a set. The aligned sets of data are shown in Supplementary Fig. S3C,D and were used for downstream analysis (Fig. 3E). Regions of end and lattice intensities were defined manually.

#### b) Analysis of guest-host experiments

Analysis of guest-host experiments was performed using a custom script written in Matlab (R2018b) including the image processing toolbox. Fluorescence microscopy images of guest/host condensates were loaded after rolling ball (50 pixel) background subtraction using ImageJ. Condensates were identified in the mCherry or Alexa-647 fluorescence channel (‘tag’). The positional information was used to quantitatively evaluate co-localization of Mal3-truncates:GFP (guest molecules, see Fig. 5B and Supplementary Fig. 4A,B) and the tag (see Supplementary Fig. S4C).

The procedure consisted of converting the mCherry/Alexa-647 image to a binary image which can be used as an image mask (im2bw function with manually optimized threshold levels ∼0.05). The image mask was then used to detect condensates and evaluate their positions, major and minor axes lengths, and the mean intensity, using the regionprops function for centroid regions.

#### c) Analysis of Mal3/Tea2/Tip1:GFP velocities

Single molecule traces of Mal3/Tea2/Tip1:GFP complexes were recorded at concentrations of 200 nM Mal3, 1 nM Tea2, and 150 nM Tip1:GFP under MPET conditions. A total number of N=148 Tip1:GFP traces remained after automated detection in seven kymographs using Kymobutler (Jakobs et al., 2019), and manual exclusion of obscure traces (crossings, merging, or tracks that reach the MT end). We calculated a median of 0.23 um/sec and a standard error of the mean of 0.06 um/sec. The velocity of Mal3/Tea2/Tip:GFP clusters in the presence of PEG35k (Figure 1H) was assessed after transfer events between microtubule ends and surrounding microtubules. We manually measured N=47 traces with a median velocity of 0.12 um/sec and a standard error of the mean of 0.018 um/sec (see Supplementary Fig. S6D).

## Acknowledgments

We are grateful to all group members of the Dogterom as well as Akhmanova (Utrecht University) labs for many discussions on microtubule end tracking proteins during ERC Synergy meetings. Initial droplet assays were performed during the Physiology Course 2017 at the MBL in Woods Hole. This work was supported by the following grants awarded to M.D.: FOM programme nr. 110 from the Netherlands Organisation for Scientific Research (L. Reese), European Research Council Synergy grant 609822 (V. Volkov), and Sinergia grant 160728 from the Swiss National Science Foundation (E. vd Sluis, R. Maan).

## Author contributions

All authors planned experiments, analyzed data, and discussed results. E.vdS. purified the proteins. R.M. performed TIRF experiments including Tea2 titration experiments. R.M. and L.R. performed droplet assays, following the initial observation by M.R.K. L.R. performed guest-host assays, theoretical modelling and analyzed data of Tea2 titration experiments. V.A.V. performed electron microscopy and 3d reconstruction with help of A.J.. W.E. coated grids with gold and contributed tilt-series acquisition scripts. N.A. contributed cryo-CARE automation scripts. V.A.V., R.M., L.R., and M.D. wrote the paper.

