## Supplementary Materials for "Multivalent interactions facilitate motor-dependent protein accumulation at growing microtubule plus ends"

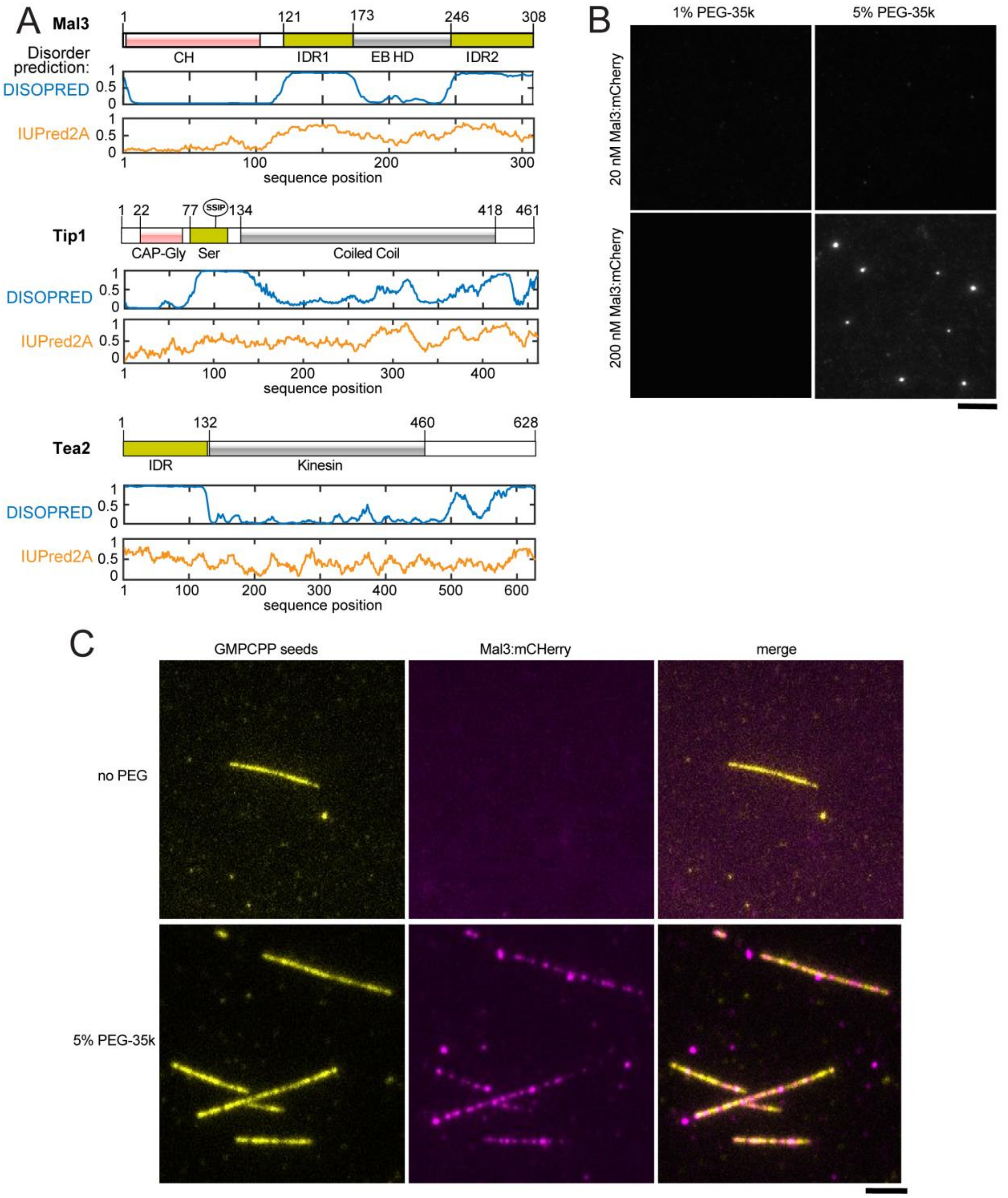


**Supplementary Figure S1. (A)** Protein regions of Mal3, Tip1 and Tea2 together with disorder prediction (DISOPRED, blue, and IUPred2A, orange). In particular, the serine-rich part of Tip1 and tails of Tea2 are predicted differently in DISOPRED3 and IUPred2A. **(B)** Titration of PEG-35k percentage and Mal3 concentration required to achieve droplet formation. **(C)** Mal3:mCherry interaction with fluorescently labelled GMPCPP seeds in the absence of PEG (top row) or in the presence of 5% PEG-35k. Scale bars: 5 μm.

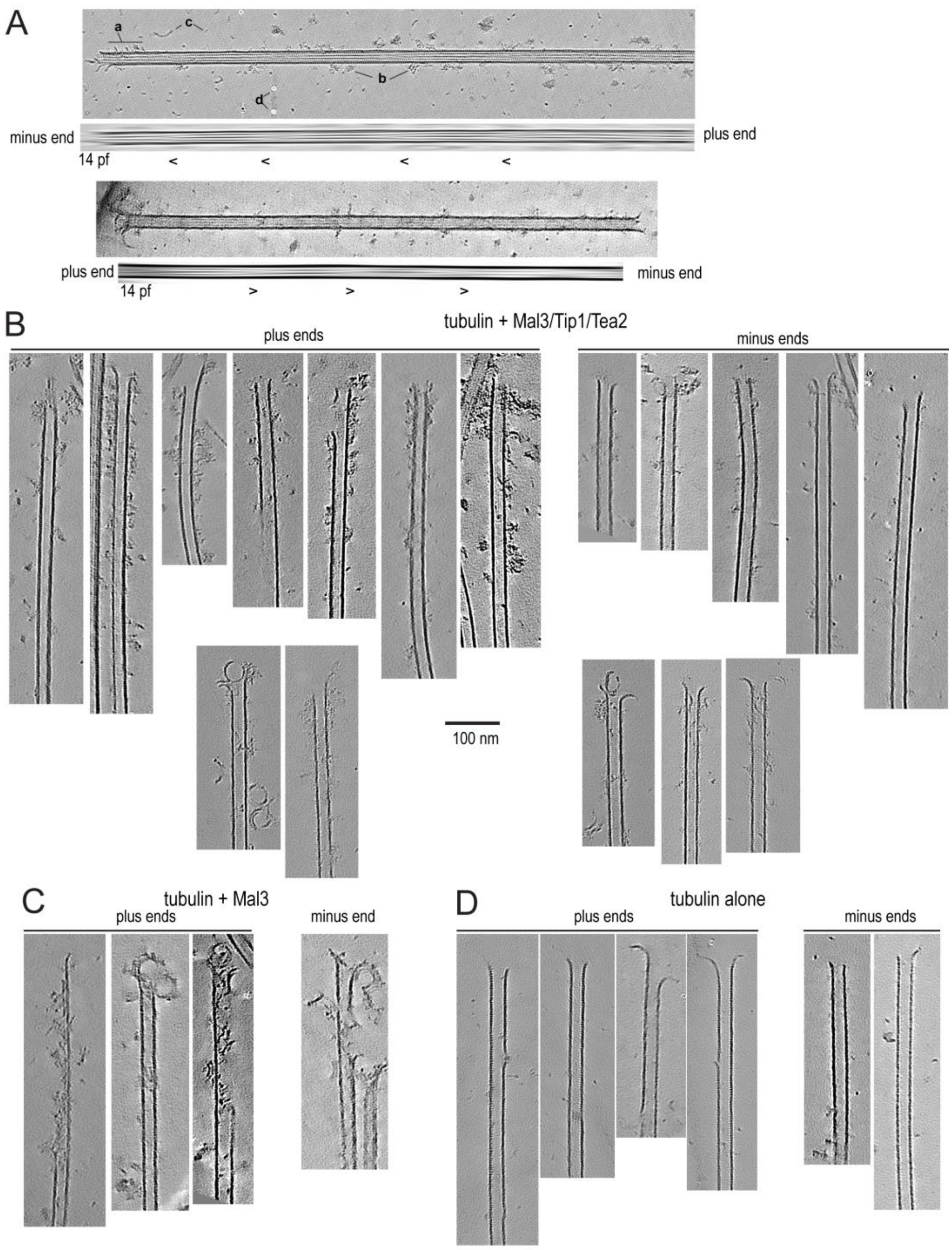


**Supplementary Figure S2. (A)** Summed tomogram slices containing microtubules (top) and Fourier-filtered subsets containing microtubule moiré pattern (bottom). Two examples are shown: a 14-protofilament microtubule with the minus end pointing up, and a 14-protofilament microtubule with the plus-end pointing up. **a** – comet at the minus-end, **b** – Mal3/Tip1/Tea2 oligomers bound to microtubule lattice, **c** – soluble tubulin oligomers, **d** – gold particles with erased densities. **(B)** Examples of plus- and minus-ends of microtubules grown in presence of Mal3, Tip1 and Tea2. Two examples of ends with unclear polarity are shown at the bottom. **(C)** Examples of plus- and minus-ends of microtubules grown in presence of Mal3. **(D)** Examples of plus- and minus-ends of microtubules grown in the absence of additional proteins. Scale bars: 100 nm.

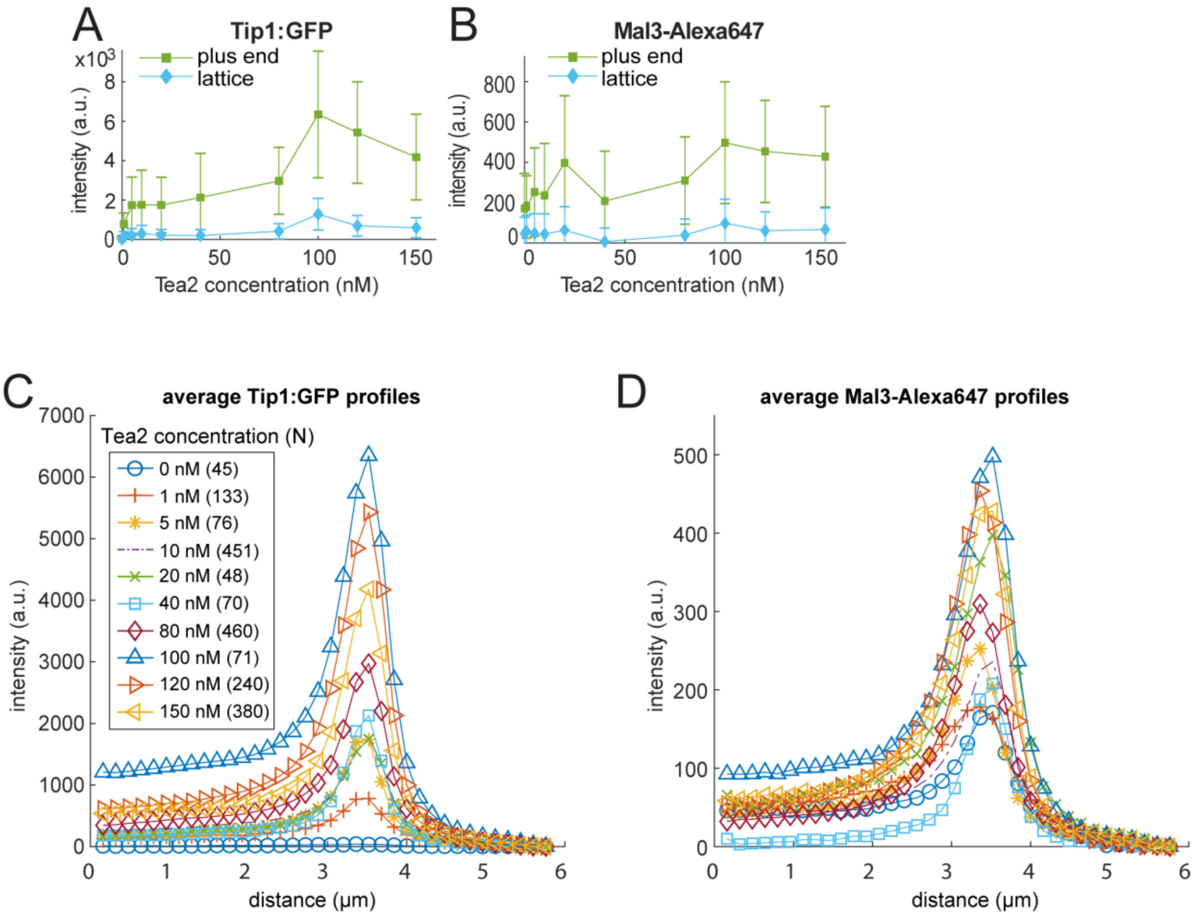


**Supplementary Figure S3. Intensity profiles of Mal3, Tip1, and Tea2 depend on motor concentration in MPET assays. ((A)** Tip1:GFP intensity on the microtubule lattice as well as on the microtubule tip strongly depends on the concentration of Tea2**. (B)** The localization of Mal3-Alexa647 is largely independent of the Tea2 concentration. **(C-D)** Intensity profiles of Tip1:GFP (C) and Mal3-647 (D) in the Tea2 titration experiments. Intensity profiles were extracted from TIRF microscopy of dynamic microtubules. (C) Tip1:GFP intensities strongly depend on the concentration of Tea2. (D) The intensities of Mal3-647 is less affected by the Tea2 concentration. The number of observed intensity profiles per condition is indicated in the legend of panel (C).

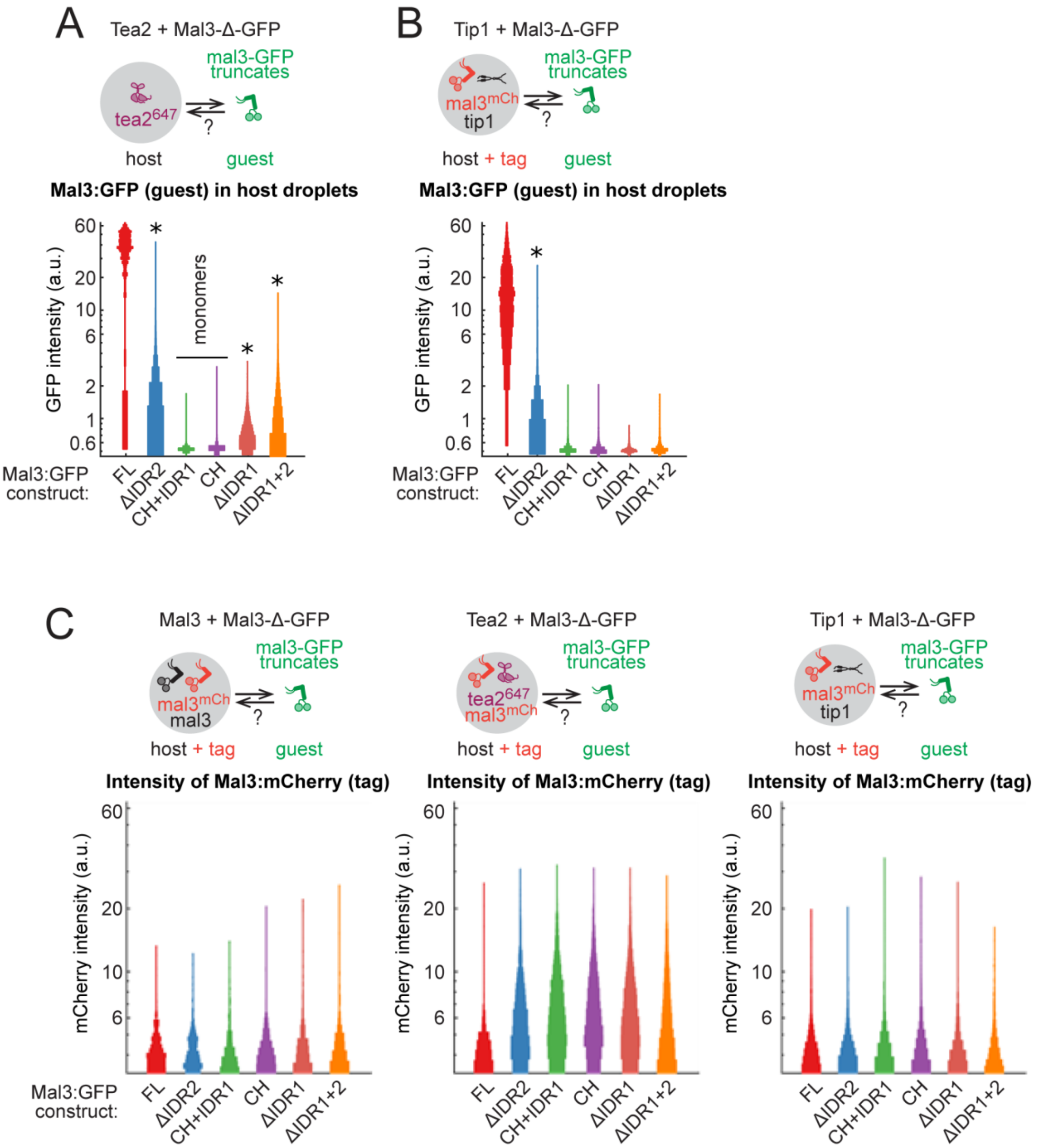


**Supplementary Figure S4. (A)** Droplets of Tea2-647 (host) were allowed to recruit Mal3:GFP constructs (2nM, guest) in presence of PEG-6k. The graph shows distribution of Mal3:GFP construct intensity in the host droplets. **(B)** Droplets of unlabelled Tip1 (host) tagged with FL-Mal3:mCherry (2 nM, tag) were allowed to recruit Mal3:GFP constructs (2nM, guest) in presence of PEG-6k. The graph shows distribution of Mal3:GFP construct intensity in the host droplets. **(C)** Intensity of Mal3:mCherry tag in experiments presented in Figure 4C (Mal3:GFP constructs recruited to unlabelled Mal3 droplets), and panels A and B.

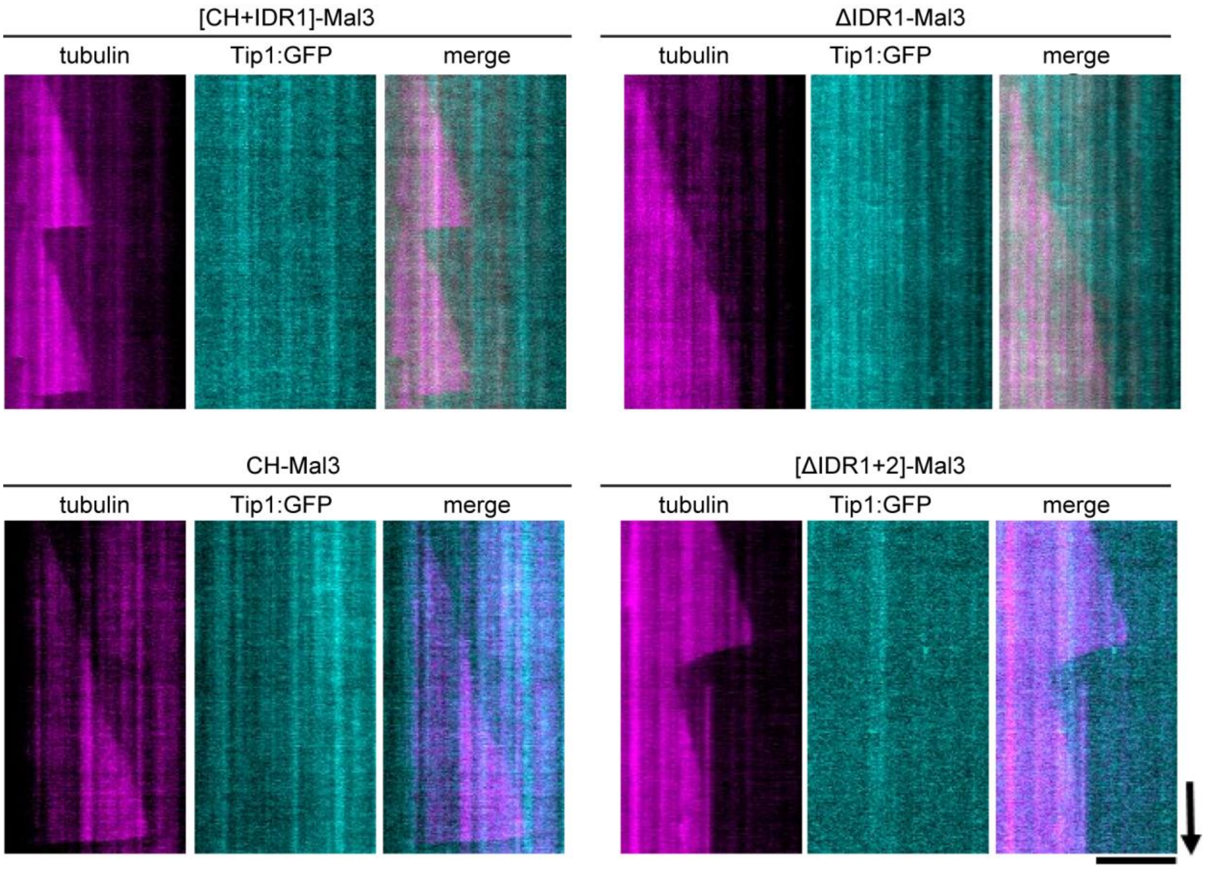


**Supplementary Figure S5.** Mal3 truncates lacking the disordered region IDR1 neither show motor transport at the lattice nor Tip:GFP accumulation at the plus end. Scale bars: 5 μm (horizontal) and 60 s (vertical).

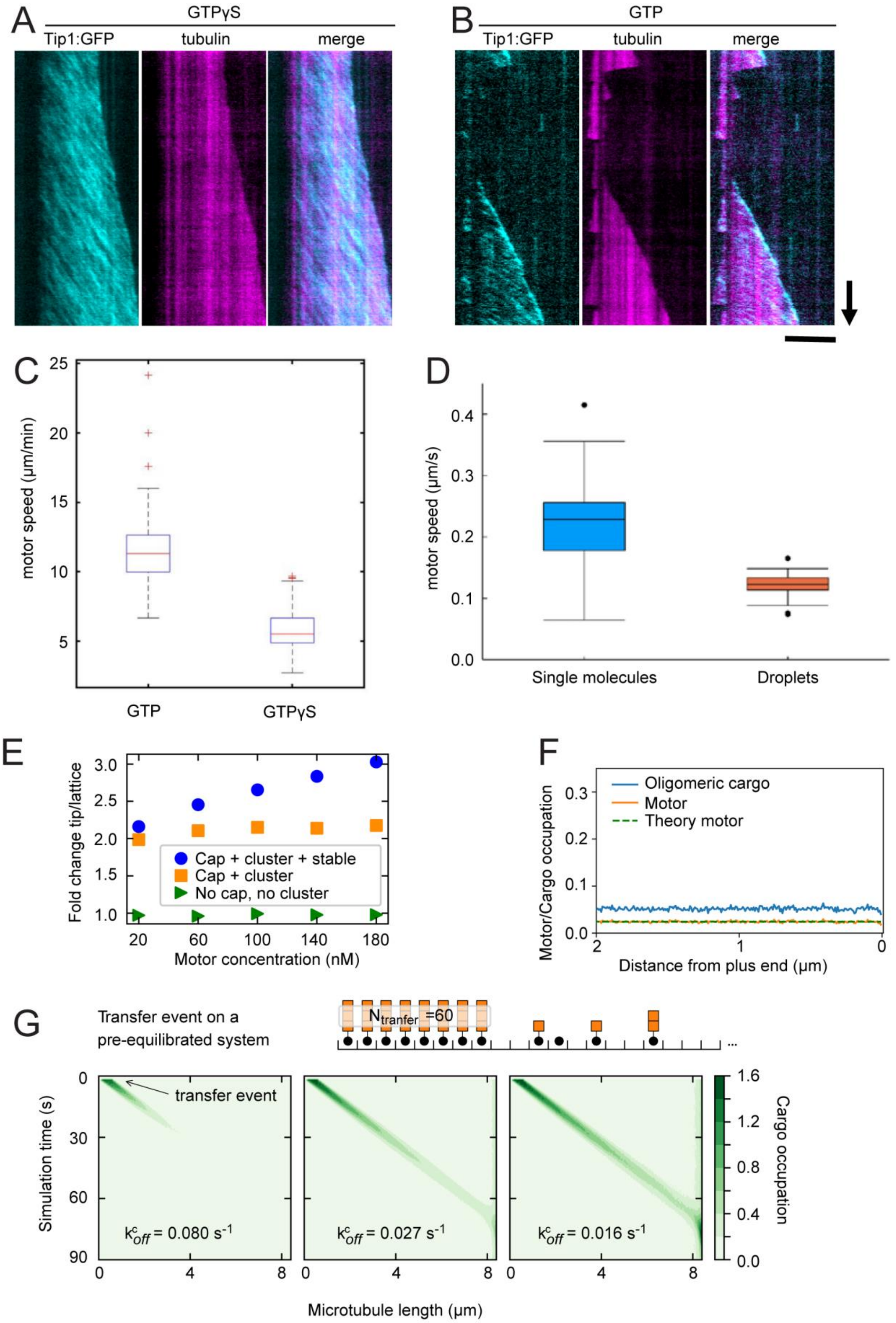

A
GTPγS
Tip1:GFP
tubulin
merge
B
GTP
Tip1:GFP
tubulin
merge
C
motor speed (µm/min)
GTP
GTPγS
D
motor speed (µm/s)
Single molecules
Droplets
E
Fold change tip/lattice
Cap + cluster + stable
Cap + cluster
No cap, no cluster
Motor concentration (nM)
F
Motor/Cargo occupation
Oligomeric cargo
Motor
Theory motor
Distance from plus end (µm)
G
Transfer event on a pre-equilibrated system
N tranfer =60
transfer event
Simulation time (s)
k^c_off = 0.080 s^-1
k^c_off = 0.027 s^-1
k^c_off = 0.016 s^-1
Microtubule length (µm)
Cargo occupation

**Supplementary Figure S6. (A and B)** MPET reconstituted in the presence of 1 mM GTP$\gamma$S (A) or 1 nM GTP (B). Scale bars: 5 μm (horizontal) and 60 s (vertical). **(C)** 100 traces were collected from 30 kymographs for both the experimental conditions to compare motor speed at GDP and GTP$\gamma$S microtubule lattice. **(D)** A comparison between the velocities of Mal3/Tea2/Tip1:GFP complexes (200 nM Mal3, 1 nM Tea2, 150 nM Tip1) and Mal3/Tea2/Tip1:GFP condensates in the presence of crowding agents (cf. Fig 1H) shows an approximate 50% reduction in plus-end directed motility (see Materials and Methods for details). We attribute this difference to the presence of the crowding agent, which effectively increases the viscosity of the surrounding solution and likely increases the cohesion of the Mal3/Tea2/Tip1:GFP condensate during molecular transport and transfers between microtubules, see Fig. 1H. **(E)** Fold change tip/lattice cargo occupation for increasing concentrations of motors, for the different models presented in Fig. 6. The model containing stable cargo clusters and motor slowdown at the cap displays a non-linear accumulation of cargo at the end, which translates into a concentration dependent increase in fold change tip/lattice. **(F)** Motor and cargo occupation in the absence of cargo clusters and motor slowdown. **(G)** The run-length of deposited comets depends on the motor dwell time in motor-cargo clusters. Motor concentration corresponds to a motor concentration of 100 nM.

**Supplementary Table S1: protofilament number and polarity of microtubule ends**

| | occurrence | polarity known |
|---|---|---|
| **12 pf** | 1 | 1 |
| **13 pf** | 8 | 0 |
| **14 pf** | 30 | 30 |
| **15 pf** | 1 | 1 |
| **total ends** | **40** | **32** |

**Supplementary Table S2: Parameters for the theoretical model**

| Parameter | Simulation | Experimental observation |
|---|---|---|
| Microtubule lattice constant | 8.4 nm | |
| Motor velocity GDP lattice | V = 20 sites/sec | 10 µm/min |
| Motor velocity GTP/GTP* lattice | $V_{cap}$ = 10 sites/sec | 5 µm/min |
| Motor landing rate | 0.00002 $site^{-1}sec^{-1}nM^{-1}$ | 0.00002 $site^{-1}sec^{-1}$ at 1 nM |
| Motor attachment rate | $k_{on}$ = [Tea2] * 0.00002 $site^{-1}sec^{-1}nM^{-1}$ | |
| Motor detachment rate | $k_{off}$ = 0.08 $sec^{-1}$ | 2 µm run-length |
| Motor detachment rate cluster | $k^{c}_{off}$ | NK |
| Microtubule growth speed | $V_{MT}$ = 6 sites/sec | 3 µm/min |
| Motor detachment lattice end | V or $V_{cap}$ | Bieling et al. |
| GTP maturation | $k_m$ = 1.4 $sec^{-1}$ | 1.4 $sec^{-1}$ (Maurer et al.) |
| GTP hydolysis | $k_h$ = 0.24 $sec^{-1}$ | 0.24 $sec^{-1}$ (Maurer et al.) |
| Cargo-motor attachment | 0.03 $site^{-1}sec^{-1}$ | NK |
| Cargo-motor detachment | 0.01 $sec^{-1}$ | NK |
| Cargo-cargo attachment | 0.03 $site^{-1}sec^{-1}$ | NK |
| Cargo-cargo detachment | 0.01 $sec^{-1}$ | NK |

| **Additional parameters for simulations** | | |
|---|---|---|
| Motor density | ρ = K/(K+1) | |
| Motor association constant | K = $k_{on}/k_{off}$ | |
| Motor entering rate minus end | α = V *ρ | |
| Simulated motor concentrations | 20, 40, 60, 80, 100, 120, 140, 160, 180 nM | |
| Simulated motor densities | 0.0050, 0.0099, 0.0148, 0.0196, 0.0244, 0.0291, 0.0338, 0.0385, 0.0431 average motor occupation probabilities | |
| Simulated effective cap size | On average half of the lattice sites have hydrolysed at ~20 sites away from the plus end. | |
| System size | N = 1000 lattice sites | L = 8.4 µm |
| Equilibration time | 100 000 sec | |
| Sampling per condition | 10 000 | |
| Sampling time | N/V = 50 sec | Time a motor needs to travel 1000 sites |

**Supplementary Video Legends**

**Video S1: Droplet fusion of full length Mal3.** Mal3 forms droplets in the presence of crowding agents. Fusion events of micron-sized droplets were observed at 24 µM Mal3 and 10% Peg 6k using DIC microscopy.

**Video S2: Droplet transfer between two microtubules.** Under crowding conditions, Mal3, Tea2 and Tip1 formed droplets at the microtubule tip. These droplets spread on the GDP microtubule lattice and fuse at the plus end. Scale bar: 5 µm.

**Video S3: MPET and fixed seed in the presence of crowding agent.** In the presence of 5% Peg 35k, Mal3 was observed interacting with the seed to support MPET on GMPCPP lattice. Biotinylated seed was made to attach the glass surface using Biotin-neutravidin binding. TIP1:GFP transported by Tea2 motors can be seen accumulating at the plus end and form a droplet that grew over time. Scale bar: 5 µm.

**Video S4: MPET and moving seed in the presence of crowding agent.** Deposition of Tip:GFP was observed from the plus end of the non-attached seed. Scale bar: 5 µm.

**Video S5-S8:** 3D rendering of denoised and segmented cryoET volumes (see Materials and Methods for details). Cyan shows microtubules and tubulin, orange – any densities not recognized as microtubules or tubulin. **Video S5**: plus-end of a microtubule grown in presence of Mal3, Tip1 and Tea2. **Video S6**: minus-end of a microtubule grown in presence of Mal3, Tip1 and Tea2. **Video S7**: plus-end of a microtubule grown in presence of Mal3 alone. **Video S8**: plus-end of a microtubule grown in presence of tubulin, without any additional proteins.
